# Systematic Elucidation and Validation of OncoProtein-Centric Molecular Interaction Maps

**DOI:** 10.1101/289538

**Authors:** Joshua Broyde, David R. Simpson, Diana Murray, Federico M. Giorgi, Alexander Lachmann, Peter K. Jackson, E. Alejandro Sweet-Cordero, Barry Honig, Andrea Califano

## Abstract

The largely incomplete and tissue-independent nature of cancer pathways represents a key limitation to the ability to elucidate mechanistic determinants of cancer phenotypes and to predict adaptive response to targeted therapy. To address these challenges, we propose replacing canonical cancer pathways with a more accurate, comprehensive, and context-specific architecture – dubbed a Protein-Centric molecular interaction Map (PC-Map) – representing modulators, effectors, and cognate binding-partners of any oncoprotein of interest. To reconstruct these complex molecular architectures *de novo*, we introduce a novel OncoSig algorithm. Validation of a lung adenocarcinoma specific (LUAD) KRAS-centric PC-Map recapitulated known KRAS biology and, more critically, identified a novel repertoire of proteins eliciting synthetic lethality in KRAS^G12D^ LUAD organoid cultures. Showing the generalizable nature of the algorithm, we elucidated PC-Maps for ten recurrently mutated oncoproteins, including KRAS, in distinct tumor contexts. This revealed a highly context-specific nature of cancer’s regulatory and signaling architectures to an unprecedented degree of resolution.

## INTRODUCTION

The idea that individual gene products may work in concert within highly conserved, mechanistic pathways, leading to coordinated activity of multiple gene products and metabolites, has long been a paradigm of molecular biology, especially in cancer research^1–3^. Recent results from the systematic reverse engineering of molecular interactions, however, challenge this view by highlighting critical limitations of canonical pathway representations^4^. In contrast to literature diagrams that depict pathways as relatively universal and mostly linear chains of events, actual molecular events in the cell are processed by a machinery that is neither universal nor linear but rather, exquisitely tissue-specific, feedback-loop rich, and too complex to be visually represented.

More importantly, key molecular interactions that mediate pathologic activity of recurrently mutated proteins and adaptive response to targeted inhibitors, within specific tumor contexts, are generally poorly represented in pathway databases, such as KEGG^5^, Gene Ontology^6^, BioCarta^7^, Reactome^8^, SPIKE^9^, Pathway Commons^10^, and Ingenuity^11^. Indeed, the interactions represented in these databases are generally supported by orthologous relationships in model organisms, noisy high-throughput assays in non-physiological contexts, or literature curation. As a result, most context-specific differences are lost, such as the differential activity of associated inhibitors in BRAF signaling between colon cancer and melanoma, leading to incorrect hypotheses about clinical utility^12^. Similarly, auto-regulatory loops and tissue-specific interactions that were not represented in canonical PI3K and MAPK pathways, were shown to be responsible for inducing adaptive responses that ultimately led to failure of otherwise promising targeted inhibitors^13, 14^.

To address such issues, we propose an integrative framework (OncoSig) for the accurate and systematic *de novo* reconstruction of context-specific, Protein-Centric molecular interaction Maps (PC-Maps). These represent the comprehensive molecular architecture necessary to support the function of a specific protein, including key oncoproteins, as implemented by three molecular interaction layers (**Figure 1a**): (a) the repertoire of upstream modulators of the Protein’s activity, such as genes harboring genetic and epigenetic variants that may contribute to its dysregulation (orange), (b) the Protein’s cognate binding partners with whom if forms stable or transient complexes (thick lines), and (c) the repertoire of downstream effectors that mediate the Protein’s pathophysiologic function (blue). Cognate binding partners may include both modulator and effector proteins. In addition, some proteins may be involved in auto-regulatory loops and thus simultaneously function as both upstream modulators and downstream effectors effectively providing a fourth functionally important interaction layer (gray lines). PC-Maps further prioritize predictions for potential synthetic lethal interactors (purple). Notably, while there has been significant focus on the reconstruction of individual transcriptional, post-transcriptional, and post-translational molecular interaction networks, development of technologies for the reconstruction of such a multi-layer, integrative logic is still elusive. This is critical because cellular phenotypes are not the results of these layers working in isolation but rather of their complex interplay.

**Figure 1.**
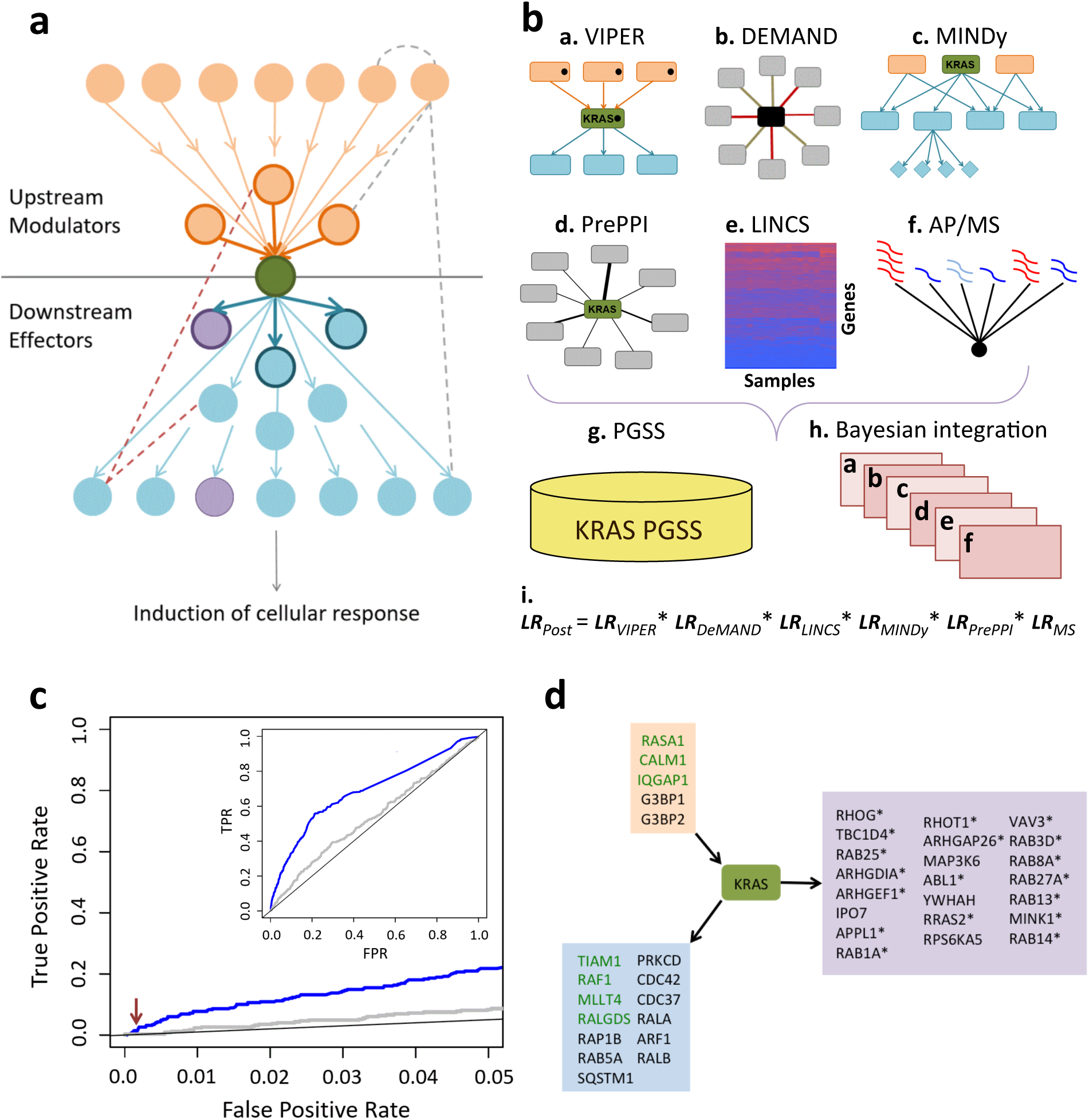
OncoSig Naïve Bayes classifier: Overview, performance, and predictions. **(a)**: A schematic of an interaction network surrounding a gene product or metabolite (green). Upstream modulators (orange) and downstream effectors (blue) interact with the gene product or other modulators via direct physical interactions (darker lines) and indirectly via transcriptional regulation or signaling cascades (lighter lines) resulting in the induction of a cellular response. Modulators and effectors may regulate other proteins in the interaction network via transcriptional regulation (pink and gray dotted lines). A subset of modulators and effectors may also be synthetic lethal (purple) with the gene product. **(b): (a)** The VIPER algorithm elucidates upstream regulators (orange) that cause a change in KRAS activity when mutated (black dots), and downstream KRAS effectors (blue) that change activity when KRAS (green) is mutated (black dot). **(b)** The DeMAND algorithm predicts dysregulated edges within a molecular interaction network. Red edges are dysregulated interactions between a protein node (black), and its partners (grey) when KRAS is mutated. **(c)** The MINDy algorithm predicts signaling molecules (orange) that co-regulate TFs (blue) with KRAS (green), leading to a change in expression of a TF’s targets (blue diamonds). (**d)** The PrePPI algorithm predicts novel KRAS (green) protein-protein interaction partners (grey). **(e)** LINCS perturbation data provides information on whether the mRNA expression levels of genes decrease (blue) or increase (red) when KRAS is knocked down. **(f)** AP-MS provides peptide counts of putative partners of the KRAS effectors RALGDS, RALA, RALB and TBK1. Blue and red indicate low and high peptide counts, respectively. **(g)** A positive gold standard set (PGSS) of proteins involved in KRAS signaling (yellow) was used to train a Naïve Bayes Classifier **(h)** that integrates the pieces of evidence **a** through **f**. **(i)** Every protein is assigned a final Likelihood Ratio (*LR*_Post_) which represents the odds (above random) of participating in KRAS-regulated signaling. **(c)**: ROC curves show the performance of the Naïve Bayes classifier (blue curve), Pearson’s correlation between mRNA expression of KRAS and mRNA expression of other proteins in LUAD (grey curve), and random performance (black curve) below a False Positive Rate threshold of 5%. The inset shows the full ROC Curve. The red arrow corresponds to a *LR*_Post_ threshold of 230. Figure 1SA provides the fraction of PGSS proteins recovers as a function of *LR*post. **(d)**: The top 40 predictions and KRAS (green box) discovered by the Naïve Bayes OncoSig Classifier. Orange and blue boxes contain, respectively, known upstream regulators and downstream effectors that are successfully recovered by the classifier. Green text indicates proteins known to interact with KRAS via a physical protein-protein interaction. The purple box shows novel NB classifier predictions tested with the RNAi negative screen; those that were experimentally found to affect cell growth in a KRAS dependent context are marked with asterisks (see Figure 2 for experimental details).

To support a realistic validation effort, we first focused specifically on assembling and validating a KRAS-centric PC-Map. KRAS is a member of the RAS family of small GTPases ^15^. KRAS is frequently mutated in cancer, especially in lung (LUAD) and colon (COAD) adenocarcinoma, where it induces a highly aggressive form of the disease, less likely to respond to targeted therapy^16–18^. We implemented the evidence-integration core of the OncoSig algorithm using either a Naïve Bayes^19^ or a Random Forest^20^ machine learning, algorithm, supporting the integration of evidence for KRAS-specific functional and physical interactions from multiple validated reverse engineering algorithms, interactomes, and databases. We found that most novel predicted KRAS PC-Map members (18 of 22) elicited synthetic lethality in 3D spheroid assays and were highly tumor-specific, while 18 additional ones were already established KRAS-related proteins. Further, PC-Maps for recurrently mutated oncoproteins and oncopathways recapitulated known interactions, as well as novel interactions, a subset of which is consistent with synthetic lethality studies by pooled RNAi-mediated silencing.

## RESULTS

In the following sections, we describe OncoSig, a novel computational technology designed to integrate multiple individual sources of evidence (henceforth *clues*), generated from both computational and experimental assays, for the inference of PC-Map proteins. To perform evidence integration, we use two machine learning algorithms supporting either direct tracing of evidence sources used to infer each interaction (Naïve-Bayes Classifier^19^) or higher-performance but lower traceability (Random Forest^20^). For the sake of clarity and without restricting the generality of the approach, we demonstrate the analytical framework by reconstructing a LUAD-specific, KRAS-centric map, assessing both established and newly predicted functional members. To show that the approach is fully generalizable to arbitrary proteins and tumor contexts, we then infer and benchmarking PC-Maps for nine additional recurrently mutated oncoproteins/oncopathways in distinct tumor contexts.

### Clues for characterization of KRAS upstream modulators, cognate binding partners, and downstream effector proteins

We used established reverse engineering algorithms to analyze the sample-matched gene expression and mutational profiles of 488 LUAD samples from The Cancer Genome Atlas (TCGA)^21^. This LUAD set comprises 326 samples with KRAS^WT^, 134 with KRAS^mut^, and 28 with no information on KRAS mutational state. VIPER, DEMAND, and MINDy analyses include transcriptional changes, under various conditions, for 1,813 proteins annotated as transcription factors (TFs), 969 proteins annotated as transcriptional cofactors (CoFs), and 3,370 proteins annotated as signaling proteins (SPs), guided by Gene Ontology (GO) classification^6^. The VIPER, DeMAND, and MINDy algorithms described below rely, in part, on ARACNe inference of regulons for KRAS and other proteins from the LUAD gene expression data^22^. Scores from the following computational and experimental sources were integrated with Naïve Bayes classification to predict members of the KRAS PC-Map, allowing easy integration of additional evidence sources, depending only on data availability, see Methods for further detail. Notably, systematic use of mutational information helps disambiguate interaction directionality, as proposed by^23^.

**1. *VIPER***^24^: VIPER is used to compute the enrichment of the transcriptional targets (*regulon*) of the oncoprotein of interest (KRAS in this case) in differentially expressed genes, with positive and negative enrichment indicating increase and decrease in KRAS activity, respectively. Statistically significant co-segregation of KRAS activity and other genes’ mutations identifies candidate KRAS upstream modulators. Candidate downstream KRAS effectors are identified as proteins whose VIPER-inferred activity co-segregates with KRAS mutations. Thus, VIPER produces clues for (a) upstream KRAS modulators as proteins harboring missense mutations that co-segregate with differential KRAS activity, and (b) downstream KRAS effectors as proteins with differential activity in KRAS^mut^ versus KRAS^WT^ samples (**Figure 1b-a**). The final VIPER scores correspond to the *p*-values associated with either clue.
**2. *DEMAND***^25^: This algorithm identifies candidate effector proteins whose molecular interactions are dysregulated in KRAS^mut^ vs. KRAS^WT^ tumor samples. Tested molecular interactions are based on an integrative reverse engineering algorithm for the reconstruction of mixed (protein-protein and transcriptional interaction) networks^25, 26^, using LUAD-specific data from TCGA. DeMAND scores are the p-values for differential dysregulation between the two groups of tumor samples (KRAS^mut^ versus KRAS^WT^) (**Figure 1b-b**).
**3. *MINDy***^27^: This clue is based on the assumption that SPs in the same pathway will modulate overlapping TF sets. The algorithm determines whether a SP is an upstream regulator or downstream effector of KRAS by evaluating the statistical significance of the overlap of the regulons of the SP and KRAS versus the overlap of regulons of the SP and other SPs. The associated score is the *p*-value of this analysis (**Figure 1b-c**).
**4. *PrePPI***^28^: This algorithm predicts KRAS binding partners among proteins in the UniProt human proteome, by combining the evaluation of structural models for the KRAS-partner interaction with other structural and non-structural clues. The associated score is the likelihood ratio (*LR*) representing the odds above random of a specific KRAS-protein interaction (**Figure 1b-d**).
**5. *LINCS***^29^: This perturbational database provides an exhaustive repository of cellular responses to drug-, shRNA-, and cDNA-mediated perturbations, as monitored by the expression of a set of 1,000 landmark genes profiled using Luminex bead assays (L1000). We used the L1000 gene expression profiles obtained from A549 KRAS^Mut^ LUAD cells with shRNA-mediated KRAS silencing to assess TFs, coFs, and SPs as potential KRAS effectors. LINCS scores represent the log fold change of expression for each gene (**Figure 1b-e**).
**6. *Affinity-purification/mass-spec assays (AP-MS)***^30^: These assays were used to characterize candidate protein-protein interactions for four established KRAS effectors─TBK1, RALGDS, RALA, and RALB─in the KRAS^Mut^ LUAD cell line A549. AP-MS scores correspond to the protein peptide count for each effector (**Figure 1b-f**). While this evidence is not systematic, it illustrates how available evidence sources, including both systematic and *ad hoc* ones, can be integrated by the methodology.

### OncoSig_NB_: Naïve Bayes classifier implementation and performance analysis

Naïve Bayes^19^ is a well-established machine learning approach for multi-class classification problems and is robust to incomplete or missing sources of data. To use this approach, raw scores for each clue (-Log *p*-values for VIPER, DeMAND, and MINDy; *LR* for PrePPI; log fold-change for LINCS; and peptide count for AP-MS) were first binned; then clue-specific *LR*s for each bin were computed as the normalized ratio of positive and negative control KRAS-interactor-proteins in a training set with the same bin value for that clue (for instance, with the same -Log *p*-value range in the MINDy analysis). For this purpose, a training set comprising a positive gold standard set (PGSS) and negative gold standard set (NGSS) must be assembled. Specifically, given a protein P, the PGSS_P_ should include established modulators, effectors, and binding partners of P, while the NGSS_P_ should include proteins that do not physically of functionally interact with it. To assemble a PGSS_KRAS_ we selected 350 proteins annotated as KRAS-pathway related in the Ingenuity Pathway Analysis Database^11, 31^ (**Supplemental File 1**). All other UniProt proteins were included in the NGSS_KRAS_ (**Figure 1b-g**), based on the assumption that the dilution resulting from inclusion of yet-unknown true positives in this set would be minimal. For each protein, then, a global posterior Likelihood Ratio (*LR*_Post_, see **Figure 1b-i**) could be calculated as the product of all individual clue-specific *LR*s (**Figure 1b-h**), with *LR* = 1 if the clue-specific score could not be computed. The normalized *LR*_Post_ represents the global posterior probability (i.e. the odds ratio) that a protein is a member of the KRAS PC-Map, with the normalization accounting for the size of the PGSS and NGSS and the expected vs. potential number of KRAS interactors. Because there are 350 and 18,901 proteins in the PGSS_KRAS_ and NGSS_KRAS_ respectively, *LR*_Post_ = 1 corresponds to a baseline probability of 1.8% (350/19251=0.018). Higher *LR*_Post_ odds ratios were normalized to this baseline *LR*_Post_ to discover the increased probability of a protein being a member of the KRAS PC-Map. *LR*_post_ and probabilities are provided in **Supplemental File 2**.

Training and testing of the classifier was first performed by 2-fold cross-validation and a receiver operating characteristic (ROC) curve was used to assess its performance (**Figure 1C**, blue). In this curve, the True Positive Rate (TPR = TP/N_PGSS_) – also called *sensitivity* – is plotted as a function of the False Positive Rate (FPR = FP/N_NGSS_) – also called specificity, – where each point corresponds to a particular *LR*_Post_ (i.e., the red arrow denotes *LR*_Post_ = 250), and TP and FP represent the numbers of predictions in the portion of the PGSS (of size N_PGSS_) and NGSS (of size N_NGSS_) not used to train the classifier, respectively. As shown in **Figure 1c**, OncoSig_NB_ achieved a TPR = 31% at FPR = 10% for *LR*_Post_ ≥ 56 (see also the black dotted vertical line in **Figure S1a**). A virtually identical performance was also achieved with Monte Carlo cross-validation (**Figure S1b**). As a comparison reference, using Pearson’s correlation between the mRNA expression of KRAS and of other proteins (gray curve) recovers only 8% of the PGSS at FPR = 10%, which is not significantly greater than random selection (black curve) (*p* = 0.178).

It is important to note that TPR and FPR in **Figures 1c and S1** only reflect established members of the KRAS pathway in the PGSS. Thus, false positives with high *LR*_Post_ values are likely to be bona fide novel KRAS pathway members. Indeed, experimental validation (next section) established that the vast majority of the highest ranking false positive predictions in **Figure 1c** were indeed novel, *bona fide* KRAS effectors and modulators. Thus, FPR as computed from prior knowledge is highly misleading and only provides a lower bound on the method’s specificity.

We chose novel predictions (included in **Supplemental File 2**) at a stringent cutoff of *LR*_Post_ ≥ 230 for further assessment and validation (red arrowhead in **Figure 1c** and red dotted vertical line in **Figure S1**). **Figure 1d** lists the top 40 predictions: 18 of the 40 predictions (orange and blue boxes) were previously established as KRAS modulators^32–34^ or KRAS effectors ^35–45^. Green text identifies proteins that were observed to physically interact with KRAS^46, 47^. Finally, the purple box highlights the remaining 22 proteins as high-confidence novel KRAS PC-Map predictions, with asterisks denoting those that were experimentally validated, as discussed in the next section.

### Experimental validation by targeted RNAi screen in primary tumor derived organoids

We expected top novel proteins in the KRAS PC-Map (*N* = 22) to be enriched for KRAS^Mut^ synthetic lethal partners and KRAS modulators. We thus performed a shRNA-based dropout screen in 3D organoid cultures derived from a LUAD KRAS^G12D/+^/p53^fl/fl^ mouse model (**Figure 2a**). Organoids in 3D cultures represent more realistic cancer models than two-dimensional monolayer cultures and provide a better representation of the *in vivo* signaling environment ^48–53^. As depicted in **Figure 2a,** primary tumor cells were infected with pools of lentiviral shRNAs and grown in 3D culture conditions for six days until organoids formed; individual cells were then dissociated and re-seeded on Day 7 to form secondary organoids ^52^. Hairpin depletion was then quantified by differential analysis of deep sequencing data from organoids at Day 6 and again at Day 12 and used to determine which hairpins inhibited organoid growth. Positive controls included TBK1 ^54^ and NUP205 ^55^, both established KRAS^Mut^ synthetic lethal partners, and the established KRAS effectors MAPK1, AKT1, RALGDS^15^, and RASA1^56^. As a negative control and to estimate the background rate of dependencies in such screens, 25 shRNAs libraries, targeting 515 genes not expected to participate in KRAS signaling, were used as a global background pooled screen (BPS). RNAi sequences are provided in **Supplemental File 3**.

**Figure 2:**
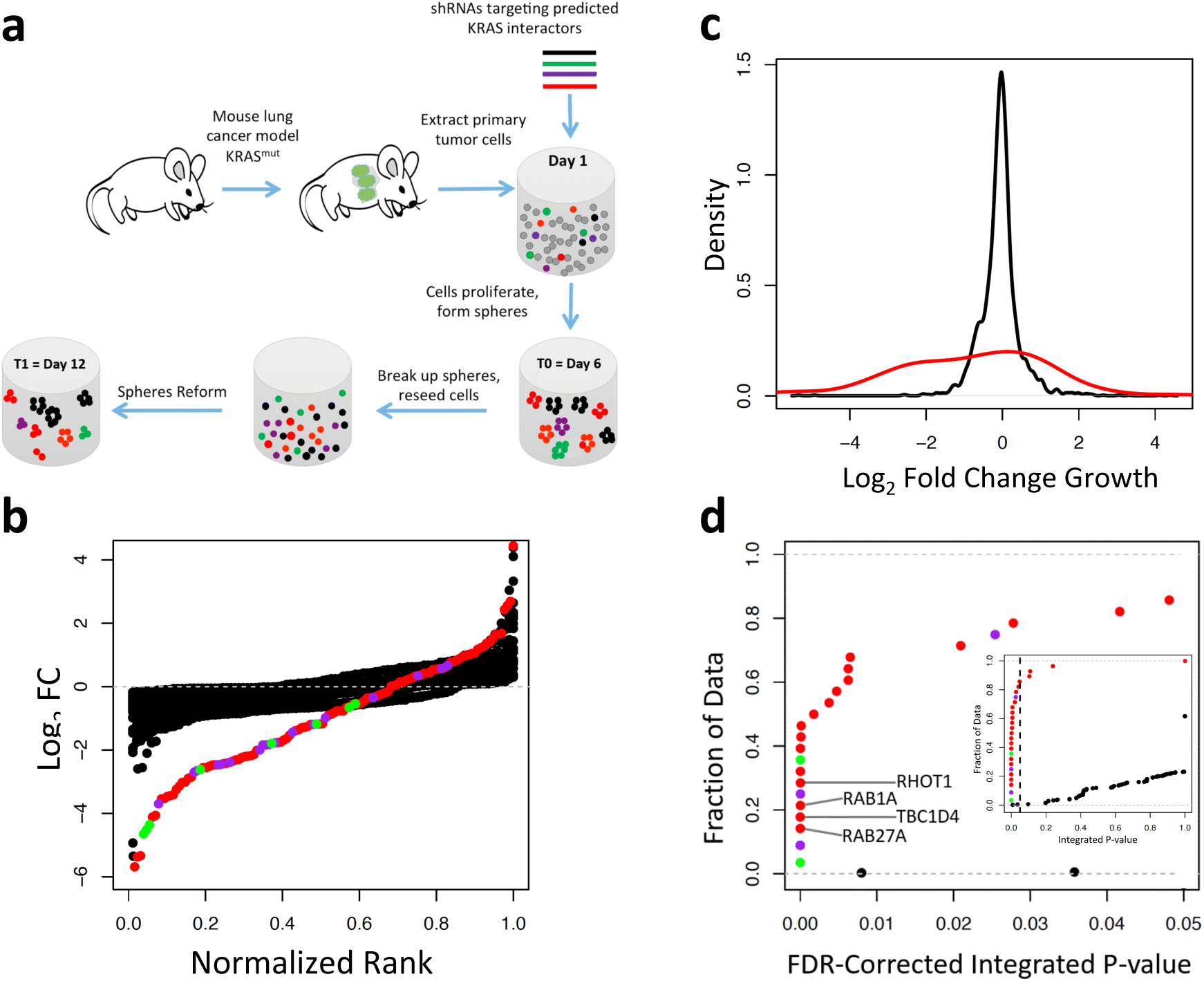
Experimental validation of OncoSig Naïve Bayes predicted KRAS functional partners. **(a)**: Schematic of the pooled shRNA negative screen experiments performed. An average of four shRNAs target each gene in the protocol implemented. KRAS^G12D/+^/p53^fl/fl^ primary tumor cells (green patches) are isolated from the mouse and placed in a semi-solid 3D matrix (cylinder). A pooled shRNA knockdown is performed (Day 1), and each cell stochastically integrates one shRNA into its DNA. Cells that integrate different shRNAs are shown as, red (representing shRNAs for novel predictions), green and purple (for positive controls, and black (for the background pool). Some cells and their daughter cells form spheroids (Day 6). The spheroids are dissociated, reseeded in a new matrix, and reform (Day 12). Fold Change (FC) of shRNA abundance is measured by deep sequencing the shRNAs at days 6 and 12. **(b)**: Plot of Log_2_FC of shRNAs targeting predicted KRAS functional partners (red), known members of the KRAS signaling pathways (RALGDS, MAPK1, RASA1 and AKT1) (purple) and two synthetic lethal positive controls (NUP205 and TBK1) (green). The black dots show Log_2_FC of shRNAs targeting 515 genes within the Background Pooled Screens (BPS) not expected to be involved in KRAS regulated signaling. The X axis is the normalized rank, calculated by ranking log_2_FC of each set of shRNAs and dividing by the number of shRNAs in that set. Each gene is represented by several dots, which correspond to different shRNAs. See Figure S2A for more details. **(c)**: Density plots of Log_2_FC for predicted KRAS functional partners (red) and an average of all BPS (black). See Figure S2B for more details. **(d)**: An empirical cumulative distribution function (eCDF) plot of the integrated p-values for the predicted KRAS functional partners, the 515 proteins in the BPS, the known members of the KRAS signaling pathways, and the synthetic lethal positive controls. The colors are the same as in panel C. The four predicted KRAS functional partners that inhibited spheroid growth to the greatest extent are labeled. The dashed vertical line in the inset indicates the p=0.05 threshold.

**Figure 2b** shows the ranked log_2_ fold change (FC) of the individual shRNA hairpins between the two time points. Consistent with previous studies^54^, growth was significantly inhibited in organoids incorporating hairpins targeting known synthetic lethal and known KRAS signaling genes (green and purple dots). Strikingly, however, a majority of hairpins targeting predicted KRAS PC-Map proteins (red dots) also inhibited organoid growth, confirming their enrichment in KRAS dependencies. In contrast, only a very small fraction of the negative control BPS hairpins (black dots) affected organoid viability. **Figure S2a** shows the log_2_ FC for the set of three to five shRNAs targeting each OncoSig_NB_ prediction.

As shown in **Figure 2c**, the average distribution of log_2_ FC values for the BPS is centered near zero and most shRNAs show less than a two-fold change (black curve; mean= ‒0.067, sd= 0.539). In contrast, the distribution for the novel predictions is highly skewed toward lower log_2_ FC values and more than a third of predicted KRAS PC-Map proteins show a decrease of four-fold or greater (red curve; mean µ = -0.852, σ = 1.952). The difference between the two distributions is highly statistically significant (*p* ≤ 2.2*10^−16^, Kolmogrov-Smirnoff test), suggesting that KRAS signaling partners predicted by OncoSig_NB_ are indeed highly enriched in synthetic lethal interactions compared to genes in the BPS negative control. **Figure S2b** shows the distributions of log_2_ FC values for each of the BPS genes. Fisher’s Method^57^ was used to integrate the one-tailed p-values for each shRNA hairpin targeting the same gene, and the Benjamini-Hochberg procedure for multiple hypothesis testing correction was performed^58^. As shown in **Figure 2d**, at an integrated *p*-value threshold of 0.05 (dotted black line in the inset plot), statistically significant inhibition of spheroid growth was observed following silencing of 18 of 22 candidate KRAS PC-Map genes (82%). In stark contrast, statistically significant inhibition of spheroid growth was observed for only 3/515 (0.58%) of BPS tested genes (*p* = 4.4×10^−16^, by Fisher’s-Exact Test, FET). Interestingly, one NB candidate RPS6KA5 (MSK1) increased spheroid growth when silenced, consistent with the discovery of generic KRAS modulators or effector (**Supplemental Figure S2c**).

We performed gene enrichment analysis of GO Biological Processes for statistically significant synthetic lethal predictions (p ≤ 0.05, by FET) using the human proteome as a null model and the Benjamini-Hochberg procedure for multiple hypothesis testing correction^58^. Consistent with expected KRAS-mediated functions, enriched GO terms fall into two main inter-related groups (**Supplemental Table 1**): GTPase-mediated signal transduction and intracellular transport. **Supplement Note 1** describes biological insight into KRAS signaling obtained from this analysis.

### Extending Naïve Bayes with a Random Forest classifier

An advantage of the NB classifier is that it supports tracing inference results to the specific clues that provide the greatest contribution to their posterior probability (*LR*_Post_, **Figure 1i**). This is useful in dissecting the biological mechanisms supporting a protein’s role in an PC-Map, including distinguishing downstream effectors vs. upstream modulators. Interpretability, however, comes at a cost because NB classifiers assume clues to be statistically independent, thus potentially under-estimating p-values when this requirement is violated. For example, the interactors of TBK1, RALGDS, RALA, and RALB, determined by AP/MS likely overlap with each other and with PrePPI-predicted KRAS interactors. Similarly, since RAB5A, for instance, is known to be regulated by RAS oncogenes^44^, proteins predicted by PrePPI to interact with KRAS are more likely to also interact with RAB5A (*ρ* = 0.47). While only PrePPI predictions for KRAS-specific protein-protein interactions were included to avoid this potential issue, other similar subtle dependencies may be harder to identify and address.

The Random Forest (RF) classifier^20^, an ensemble-based decision-tree method, is an alternative machine learning approach for integrating large-scale genomic and network interaction data. It generally outperforms the NB classifier in a wide range of problems^59^. RF classifiers are less affected by correlated clues and better at learning from a large number of relatively weak clues, thus allowing incorporation of additional information. However, they are less effective in terms of supporting the ability to trace their inferences to the specific contributing evidence sources. We thus also tested PC-Map inference using a RF classifier (OncoSig_RF_), with the additional advantage that this methodology can dissect PC-Maps seeded on one or more oncoproteins, including on a handful of key proteins in an established pathway, such as HIPPO or WNT^60, 61^. We call the proteins used to “seed” the PC-Map *Core Proteins*.

### OncoSig_RF_: Random Forest classifier implementation and performance analysis

As shown in **Figure 3a**, the RF classifier incorporates clues similar to those discussed above (**Figure 1a**) with the following differences: (a) We eliminated the AP-MS, LINCS, and DEMAND components because they were not available for all tested proteins/pathways; (b) We included ARACNe ^22^ (**Figure 3a-a**) to assess context-specific transcriptional targets of TFs, CoFs and SPs; (c) MINDy was replaced by CINDy^62^, a more recent version of the algorithm with improved sensitivity; and (d) an updated version of the PrePPI^63^ database with higher recall was used, including ∼16,000 PPIs, rather than only KRAS-specific interactions. As before, 1,813 TFs, 969 CoFs, and 3,370 (SPs) were analyzed by the regulatory network components (**Figure 3a, a-c**).

**Figure 3:**
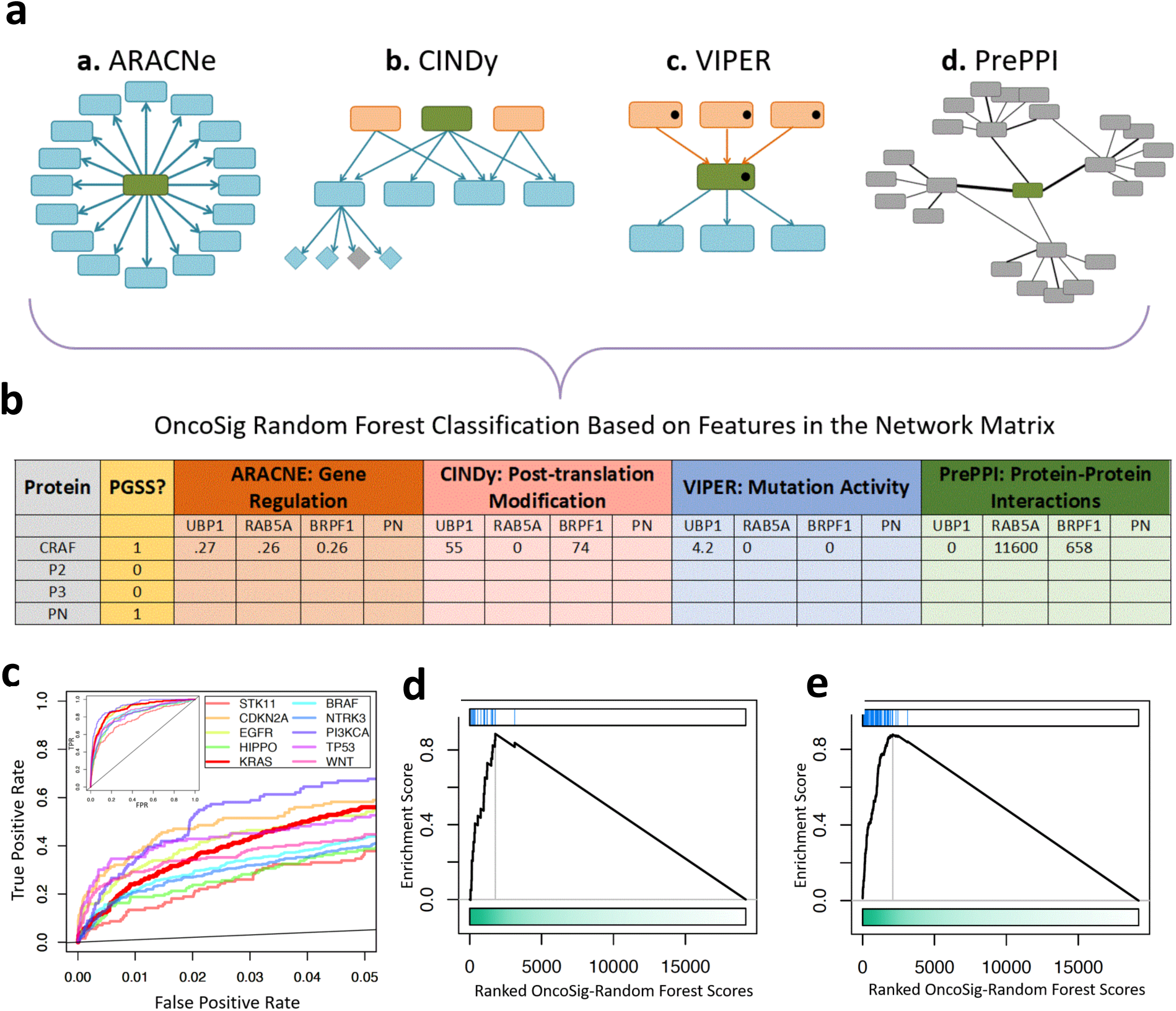
OncoSig Random Forest classifier: Overview, performance, and prediction. **(a)**: Networks used to train the OncoSig Random Forest classifier: (a) The ARACNe algorithm predicts transcription factors or signaling molecules (green) that transcriptionally regulate target genes (blue). (b) CINDy predicts signaling molecules (orange/green) that post-translationally modify transcription factors (blue boxes), which in turn leads to differential expression of a transcription factor’s targets (blue or gray diamonds). (c) The VIPER algorithm infers downstream effectors (blue) and upstream regulators (orange) for a given protein (green). VIPER associates 1) the protein (green) with a missense mutation (black dot) with the activity change of transcription factors (blue) and 2) signaling molecules (orange) with missense mutations (black dots) with activity of the protein (green). (d) PrePPI predicts interactions between a protein (green), and its functional interactors (gray). **(b)**: The networks are encoded as a matrix of feature vectors and colored as: ARACNe (orange), CINDy (pink), VIPER (blue), and PrePPI (green). The gold column corresponds to whether a protein is a member of a particular pathway’s PGSS (1) or NGSS (0). An example of a feature vector for one protein is shown for CRAF with its corresponding values. **(c)**: ROC curves depict OncoSig performance for the 10 pathways in the key below FPR = 0.05. The inset shows the full ROC curves. The thick red line represents performance for the KRAS PC-Map. **(d)**: Gene Set Enrichment Analysis (GSEA) of the 22 OncoSig Naïve Bayes predicted KRAS functional partners tested in the knockdown experiments (blue lines) and OncoSig Random Forest results for KRAS, where the ranking is based on OncoSig RF score. **(e)**: GSEA of the top 100 NB predictions (blue lines) and RF results for KRAS, where the ranking is based on OncoSig RF score.

We used OncoSig_RF_ to infer PC-Maps centered on eight of the most recurrently mutated oncogenes/tumor-suppressors (BRAF, CDKN2A, EGFR, KRAS, NTRK3, PI3KCA, STK11, and TP53)^64^ and two cancer-related pathways (HIPPO and WNT) (Harvey et al., 2013; Duchartre et al., 2016). To standardize the process, independent PGSSs for each protein/pathway were derived from the mSigDB C2 curated gene set^65^ and supplemented by KEGG^5^ (**Supplemental File 1,** columns 2-11). In each case, the NGSS comprised the UniProt human proteome with the appropriate PGSS removed.

Taken together, the clues used by OncoSig_RF_ include 2,504,215 computationally and, in many cases, experimentally supported candidate functional and physical interactions between 19,548 proteins (**Figure 3a**). These clues are encoded as a single matrix with 19,548 rows (the UniProt human proteome) and 48,931 columns (all genes products whose functional or physical interaction with a protein is supported by at least one clue). **Figure 3b**, for instance, provides quantitative feature values for the interactions between CRAF and three other gene products (UBP1, RAB5A, and BRPF1): (a) mutual information for ARACNe (orange), (b) the number of SP-CoTF/TF-gene triplets sharing statistically significant CINDy conditional mutual information (pink), (c) -log(*p*-value) for VIPER (blue), and (d) likelihood ratio for PrePPI (green), where *LR*_PrePPI_ was taken from the PrePPI algorithm after removing GO contributions. The gold-standard column designates which proteins belong to a PGSS (“1”) or to a NGSS (“0”).

The classifier was evaluated by Monte Carlo cross-validation, using the full feature matrix to produce the ten PC-Maps. Monte-Carlo cross validation ensures that predictions for each protein are generated only with *forests* that exclude *core proteins* from the training sets. An OncoSig_RF_ score (S_RF_) of 0.5 means that 50% of the trees supported a protein’s association with PC-Map core proteins (see Methods). Proteins with S_RF_ ≥ 0.5 are thus considered reliable candidate PC-Map members.

### Performance analysis of oncogene-centric pathway reconstruction

The ROC curves in **Figure 3c** assess OncoSig_RF_’s performance on the oncoproteins/oncopathways (**Supplemental File 1**). Performance improvement over random classification (black line) and gene expression correlation (not shown) is highly statistically significant (p <10^−10^ in all cases). As shown, classifier performance varies widely: For instance, at FDR = 1%, 35%-37% of established PI3K-, TP53-, and CDKN2A-pathway members were recovered (purple, pink, and orange curves), whereas 12%-20% of established STK11-, HIPPO-, and NTRK3-pathway members were recovered (salmon, green, and blue curves). Predictions for each of the ten PC-Maps are provided in **Supplemental File 4**.

### Random Forest and Naïve Bayes classifiers produce consistent predictions

The bold red ROC curve in **Figure 3d** represents the KRAS-specific OncoSig_NB_ performance. While OncoSig_RF_ significantly outperformed OncoSig_NB_, their predictions were highly consistent based on Gene Set Enrichment Analysis (GSEA)^66^ using either the top 22 OncoSig_NB_ predictions that were experimentally tested (NES = 5.4, p = 5.6 × 10^−8^) and the top 100 OncoSig_NB_ predictions overall (NES = 9, p = 1.7 × 10^−19^) (**Figures 3d and 3e**, respectively). **Figure S3** shows that the OncoSig_RF_ and OncoSig_NB_-inferred PC-Maps are similarly highly consistent when OncoSig_RF_ is trained with the Ingenuity PGSS (see **Supplemental File 1,** columns A and F).

### Performance of the KRAS-specific OncoSig_RF_ analysis

KRAS-specific OncoSig_RF_ analysis (**bold red line; Figure 3c**) recovered 61/250 (24%) and 140/250 (56%) of PGSS_KRAS_ at a FPR = 1% and 5%, respectively. Since predictions were LUAD-specific, whereas the PGSS_KRAS_ is essentially context-free, these represent high recall rates. Further, at FPR of 1% and 5%, OncoSig_RF_ makes 193 and 965 novel predictions. However, as discussed above, the ROC analysis aims only to assess the ability of the classifier to recover the PGSS, and FPR cutoffs correspond to upper bounds and result in lower apparent performance. We consider novel predictions as proteins that do not appear in the PGSS and have S_RF_ ≥ 0.5 (See **Supplemental File 4** for a list of all predictions.)

A few potential sources of performance bias are considered. First, proteins with high connectivity (i.e. hub proteins) may be favored by the analysis due to their potentially greater influence in the PC-Map of interest. However, as shown in **Figure S4**, ranking by node degree (**brown curve**) and training on a randomly permuted network (**tan curve**) recover 5-fold fewer PGSS proteins than OncoSig_RF_ (**red curve**). Second, several established KRAS modulators and effectors in the PGSS have high sequence similarity (e.g. members of the MAPK family), which may bias predictions towards sequence-similar neighbors. We generated a sequence non-redundant PGSS with CD-HIT ^67^ to cluster PGSS proteins with sequence identity ≥80%. As shown in **Figure S4b**, ROC curves using the full (**red curve**) or non-redundant (**green curve**) PGSS are similar, i.e. same recovery at FPR = 0.5% and < 1.5-fold greater recovery for the non-redundant PGSS at FRP = 1%, indicating that OncoSig_RF_ predictions are minimally affected by PGSS sequence redundancy. Finally, OncoSig_RF_ predictions are biologically relevant: They recapitulate 14% of KRAS pathways members from STRING^68^ and/or HumanNet^69^, which are widely used, curated resources for protein functional interactions (**Figure S5**). In addition, the algorithm provides many unique predictions, thus expanding the repertoire of possible KRAS signaling partners.

### The KRAS PC-Map is highly enriched in proteins eliciting synthetic lethality in KRAS^Mut^ cells

Several studies have identified synthetic lethal dependencies in oncogenic KRAS^Mut^ cells^54, 70, 71^. To check the enrichment of the KRAS PC-Map in experimentally identified synthetic-lethal proteins, the analysis was performed by removing the proteins identified by the corresponding synthetic-lethal screen from the training set. 204 genes from 19 cancer cell lines were identified as KRAS^Mut^ but not KRAS^WT^ essential based on silencing by at least one shRNA hairpin ^54^.

According to stricter criteria, 45 of these genes qualified as high-confidence KRAS^Mut^ synthetic lethal partners ^54^. As shown in **Figure 4a**, OncoSig_RF_ predictions are highly enriched in the set of 204 genes (NES = 7.73, p = 2.4×10^−14^). There is also high enrichment in the set of 45 strict synthetic-lethals (NES = 4.2, p = 1×10^−5^; **Figure S6a**). These results, along with the high rate of experimental validation for OncoSig_NB_ predictions (**Figure 3b**) and their strong enrichment in OncoSig_RF_ predictions (**Figure 3d and e**), strongly support the algorithm’s ability to identify effector proteins eliciting synthetic lethality with KRAS^Mut^.

**Figure 4:**
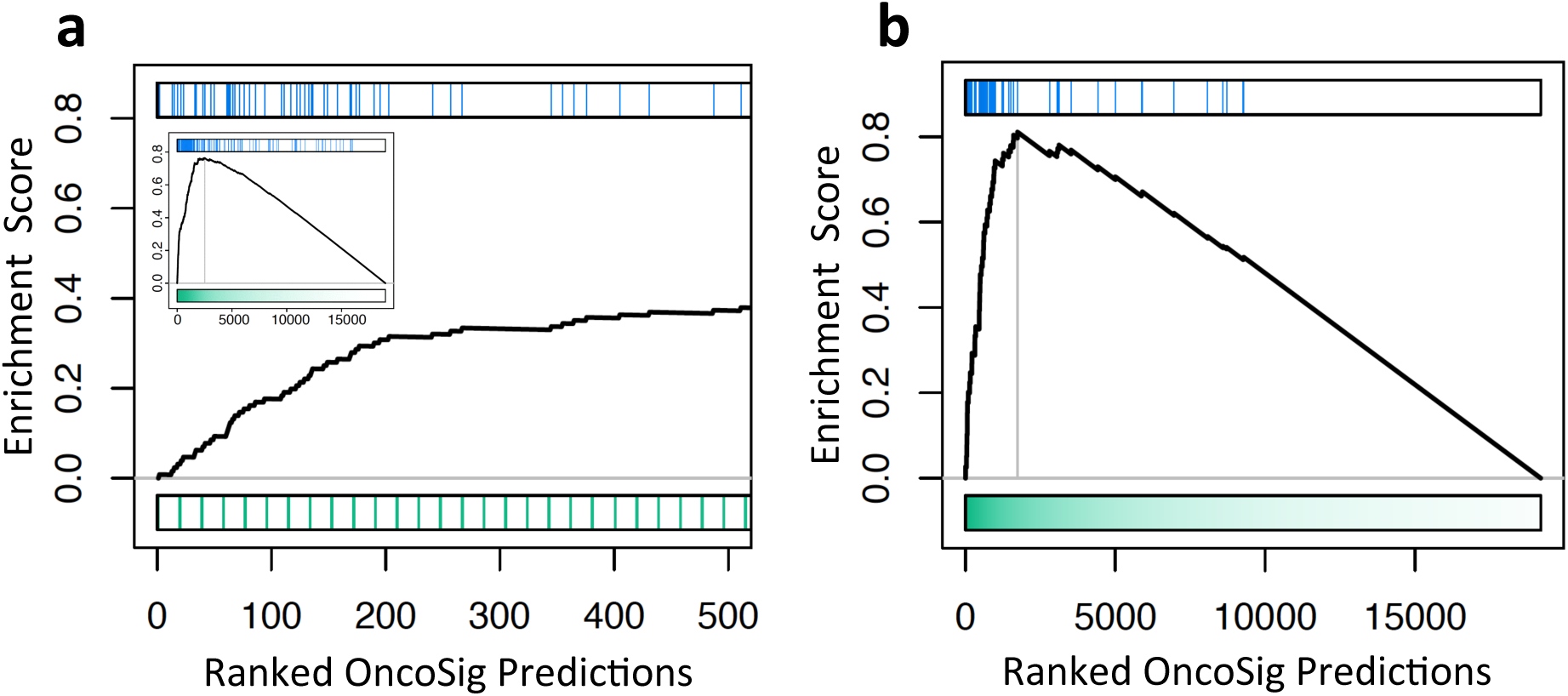
OncoSig Predictions are enriched for KRAS and EGFR synthetic lethal interactions. **(a)**: GSEA of KRAS synthetic lethal partners^54^(blue lines) and the top 500 KRAS OncoSig predictions obtained by training on a modified PGSS for which the intersection with the Barbie et al. set was removed. Inset is the GSEA using all OncoSig predictions obtained in this way, where the ranking is OncoSig score. GSEA plots for additional literature-derived sets are provided in **Figure S6**. **(b)**: Enrichment of EGFR synthetic lethal partners in the presence of an EGFR inhibitor^72^ with OncoSig predictions obtained by training on a modified PGSS for which the intersection with the Astsaturov et al. set was removed. GSEA plots for additional literature-derived sets are provided in **Figure S7**.

Predictions were also enriched in gene sets from a variety of additional KRAS-related studies, including: (a) genes eliciting synthetic lethality in KRAS^Mut^ cells treated with a MEK inhibitor ^70^ (**Figure S6b**); (b) genes contributing to ERK inhibitor resistance in KRAS^Mut^ cells ^71^ (**Figure S6c**); and (c) genes inducing oxidative stress proteins lethal to KRAS^Mut^ cells ^6, 72^ (**Figure S6d**).

To extend this type of validation to other oncogenes, we tested the algorithm’s ability to recapitulate EGFR pathway members^73^. As shown in **Figure 4c**, Predictions were highly enriched in 58 genes whose knockdown sensitized cells to EGFR-targeted inhibitors (NES = 6.0, p-value = 1.4 × 10^−9^). As shown in **Figure S7a**, Predictions were also highly enriched in the full set of ∼600 curated EGFR pathway members targeted by ^73^ (NES = 14.3, p = 2.3 × 10^−43^) and further differentiated genes that sensitize cells to EGFR-targeting drugs versus genes that do not (p = 2.0 × 10^−4^, Welch’s two sample t-test; **Figure S7b**).

### Cancer context specificity

Thus far, the evidence that was integrated by OncoSig_RF_ (**Figure 3**) was produced from TCGA LUAD cohort specific data. To assess context specificity of the predictions, we compared the LUAD-specific analysis, with equivalent analyses based on 434 TCGA colon adenocarcinoma (COAD) samples and 482 TCGA lung squamous cell carcinoma (LUSC) – the latter representing a cancer histologically more similar to LUAD than COAD. These data were used for ARACNe (**Figure 3a-a**), CINDy (**Figure 3a-b**), and VIPER (**Figure 3a-c**) analyses. The PrePPI network is independent of cancer type.

OncoSig_RF_ was bootstrapped 100 times to produce LUAD-, LUSC-, and COAD-specific KRAS-centric PC-Map predictons, using the PGSS_KRAS_. **Figures 5a-5c** are scatterplots of scores (S_RF_) for two LUAD (**5a**), one LUAD and one LUSC (**5B**), and one LUAD and one COAD bootstraps (**5c**). Only predictions with S_RF_ ≥ 0.5 in at least one contexts were considered, to eliminate irrelevant low-confidence predictions (gray dots in **Figures 5a-5c**). For LUAD-LUAD (**5a**), there are essentially no “off-diagonal” points and R^2^ = 0.99. As expected, the spread of off-diagonal predictions in the scatterplots suggests that KRAS-specific PC-Map conservation is much more significant in LUAD vs. LUSC (**5B**; R^2^ = 0.35, p = 2 x10^−267^) than in LUAD versus COAD (**7C**; R^2^ = 0.10, p = 6 x10^−165^). This shows that context-specific PC-Map differences cannot be discounted as the effect of distinct dataset analyses.

**Figure 5:**
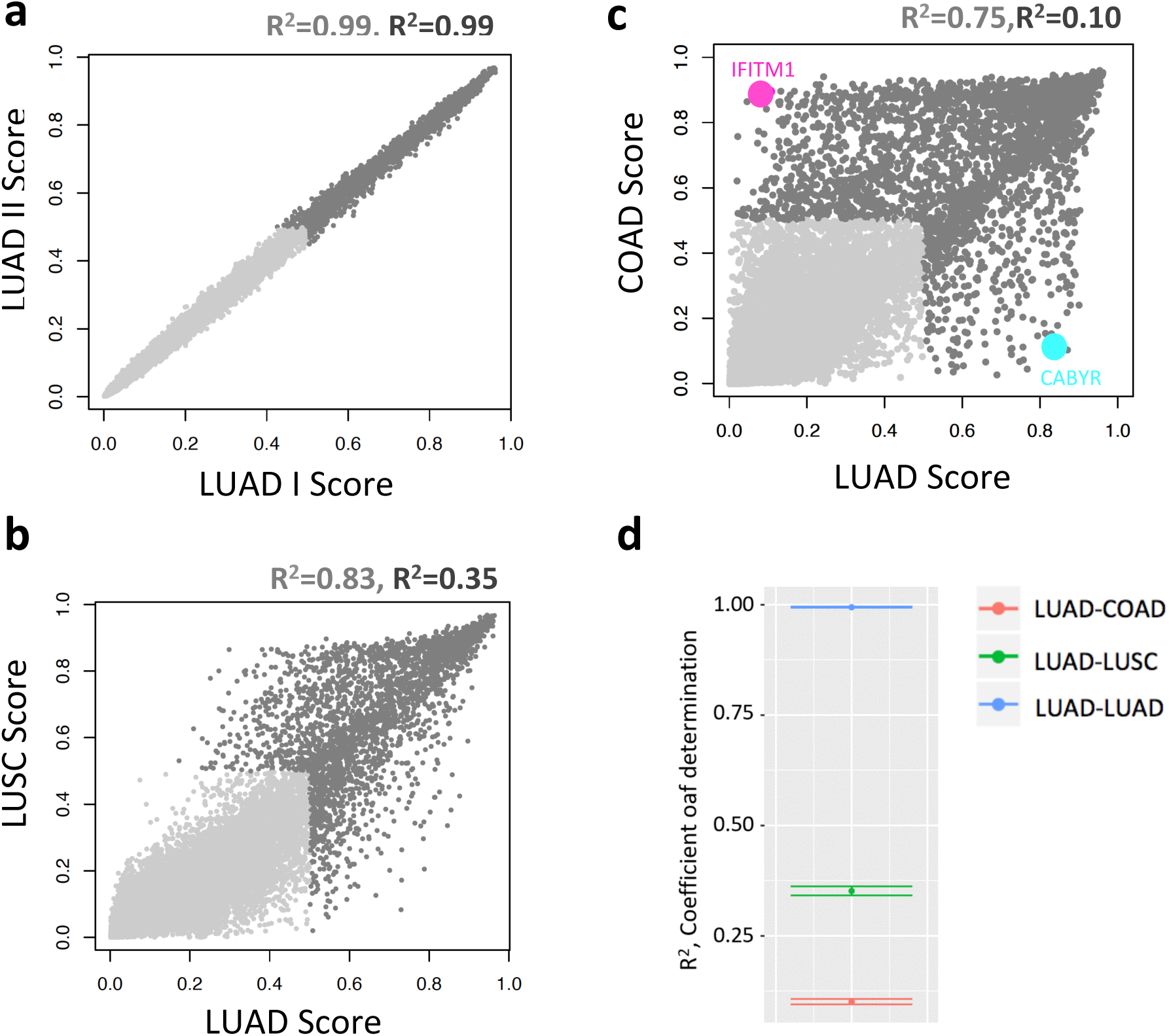
OncoSig predictions based on LUAD-, LUSC-, and COAD-derived networks. **(a)**: Plot of OncoSig scores for two OncoSig LUAD replicates. Each dot represents the scores for one protein. The light gray dots are predictions that are below 0.5 in both LUAD replicates. The light and dark R^2^ are, respectively, the squared Pearson correlation coefficient for all points, and just for the dark points. **(b)**: Plot of OncoSig scores for one OncoSig LUAD run (X axis) versus one OngoSig LUSC run (Y axis). Each dot represents the scores for one protein. The color scheme is the same as **A**. **(c)**: Plot of OncoSig scores for one OncoSig LUAD run (X axis) versus one OngoSig COAD run (Y axis). Each dot represents the scores for one protein. The color scheme is the same as **A**. The magenta and cyan points are scores for CABYR and IFITM1, respectively). **(d)**: Box plot of LUAD-LUAD (blue), LUAD-COAD (red) and LUAD-LUSC (green) squared correlation coefficients (R^2^) based on 100 OncoSig runs. Mean R^2^ (points) were calculated by taking the average R^2^ for all 100×100 runs. The bars show the standard deviation. Only predictions with a score of 0.5 or greater (e.g. corresponding to dark points in **A** and **B**) were used to calculate R^2^.

**Figure 5d** summarizes the mean R^2^ values (dots, with standard deviations denoted by bars), calculated as the average R^2^ for all 100×100 bootstraps, for the three comparisons. As expected, R^2^ = 0.99 (σ = 0.0005) for LUAD-LUAD (blue). R^2^ for LUAD-LUSC (green) and LUAD-COAD (red) are significantly lower: R^2^ = 0.35 (σ = 0.011) and R^2^ = 0.10 (σ = 0.006), respectively. Thus, LUAD OncoSig_RF_ predictions account for 35% of the variation of LUSC predictions and only 10% of the variation of COAD predictions, consistent with the decrease in histological similarity in LUSC vs. COAD.

As seen in the LUAD-COAD comparison (**Figure 5c**), many predictions have a high OncoSig_RF_ score in one context and a low OncoSig score in the other. For instance, IFITM1 (magenta) and CABYR (cyan) Overall, the KRAS-specific PC-Maps in LUAD and COAD (e.g. gray and black dots in **Figure 5c**) share 1,829 predicted interactions of which 164 are predicted to be physical, while 411 and 752 predictions are unique to LUAD and COAD, respectively (**Supplemental File 4**).

## DISCUSSION

The term “pathway” is one of the most loosely defined biological concepts. And yet, it is also one of the most frequently used in the literature. Here, we propose to replace the traditional but largely qualitative concept of “pathway” with that of the regulatory machinery (i.e., the set of gene-products and associated regulatory topology) that is necessary for a specific protein to perform its physiologic function, as well as to induce pathologic behavior when dysregulated by deleterious endogenous or exogenous perturbations. We use the term Protein-Centric molecular interactions Map (PC-Map) to model the resulting architecture. Key differences between PC-Maps and traditional pathways include (a) their systematic and principled construction (**Figure 3**), (b) their tissue-specific nature (**Figure 5**), (c) the depiction of gene-products representing upstream modulators, downstream effectors, binding partners, and members of autoregulatory loops (**Figure 6**), (d) the specific metrics associated with these roles (**Figure 6 and Supplemental File 5**), (e) their prioritization of synthetic-lethal and drug-sensitizing partners (**Figure 4**), and (f) the ability of the algorithm to seamlessly integrate many types of data and interactomes beyond those described in **Figure 3**. As such, PC-Maps provide more powerful unbiased representations than most network approaches and a more meaningful description of the regulatory networks that determine the function of a given protein.

**Figure 6:**
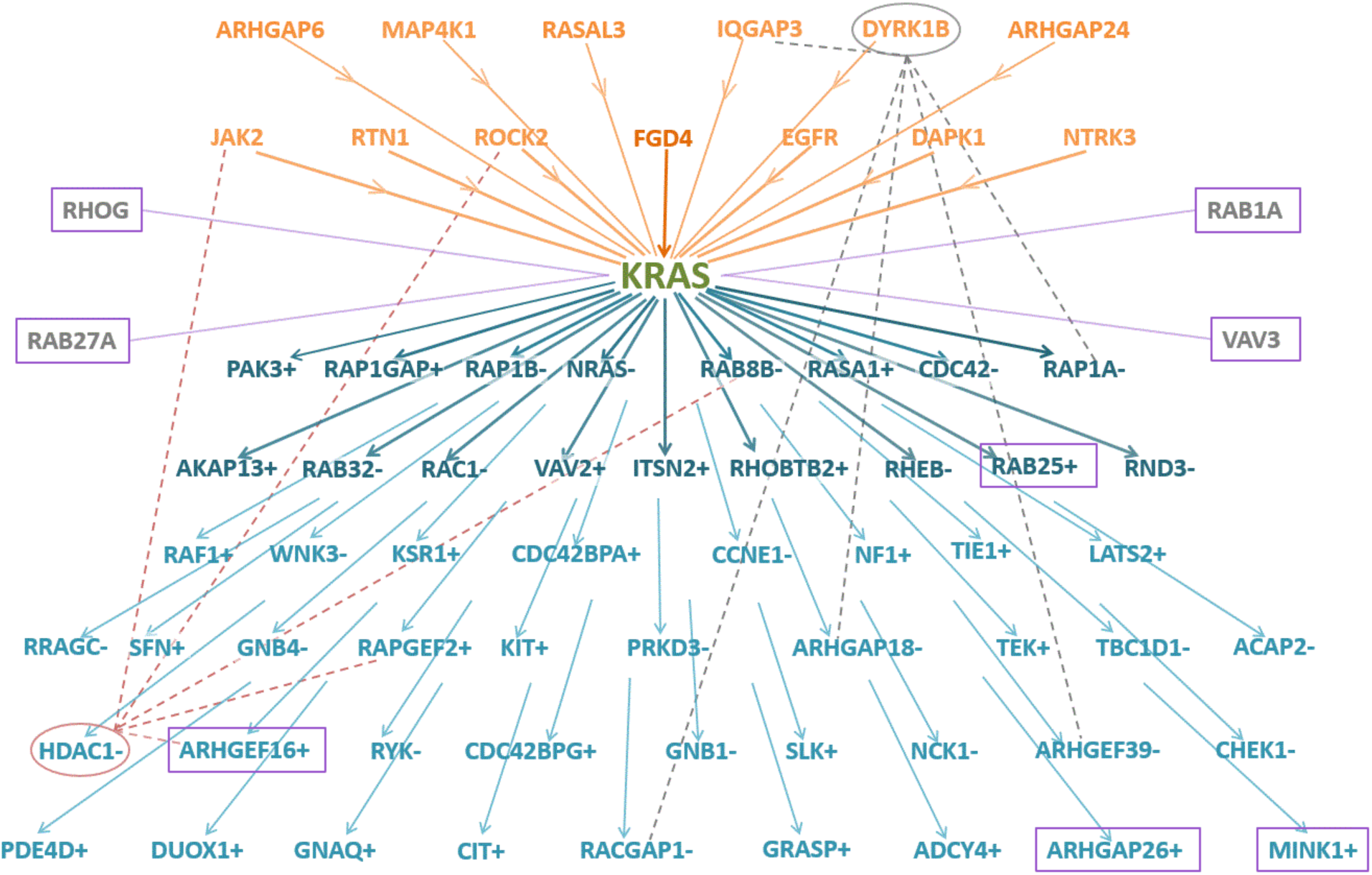
An Integrated view of KRAS LUAD PC-Map. Proteins that have an OncoSig KRAS pathway score ≥ 0.85 and a statistically significant VIPER mutation-activity interaction with KRAS in LUAD TCGA tumor samples are shown. Orange nodes indicate proteins that when mutated change KRAS activity (orange arrows, upstream regulators). Blue nodes indicate proteins that change activity when KRAS is mutated (blue arrows, downstream effectors), and + and – signs indicate whether the activity of these proteins increases or decreases, respectively. Darkened text and arrows indicate proteins predicted by PrePPI to bind KRAS in a physical PPI. Purple boxes denote experimentally tested predictions supported in the synthetic lethality assay at a p-value threshold of ≤ 0.05. Grey and rose ovals and dotted arrows indicate CINDy predicted transcription factor modulation by signaling molecules. The threshold for VIPER and CINDy interactions are the same as those used to generate OncoSig predictions, but the PrePPI predictions must have a final score *LR*_PrePPI_ ≥ 600 as well as a structural modeling score *LR*_SM_ ≥ 100, indicating further evidence of physical PPI. Note that the proteins depicted here do not correspond to the very maximum proteins predicted by OncoSig (Supplemental File 3) because, to avoid cluttering the image, only those predictions that are predicted by OncoSig and VIPER as well are shown.

PC-Maps thus encapsulate the complex biology of signal transduction as evidenced by the high rate of experimental validation of novel predictions (18/22, 82%) (**Figures 1 and 2**). Strikingly, 36 of the top 40 predicted KRAS functional partners (90%) were validated as *bona fide* members of the KRAS PC-Map based on literature and experimental validation assays. Further, as illustrated in **Figure 6**, The analysis successfully identifies EGFR and NTRK3 as upstream KRAS modulators^15, 74^ and RAF1 and CDC42 as downstream effectors^37, 75^. However, it fails to recapitulate the specific directionality of some KRAS interactions such as those with RASA1 and NF1. Similarly, a number of established direct physical KRAS interactors are successfully predicted by the analysis, such as RASA1, yet some are missed, such as RAF1. The latter may depend on critical clues not included and it is possible that the interactions may not be germane to the lung cancer context in which the analysis was performed, It is also possible that prediction efficacy is limited by the reliance on a positive gold standard set (PGSS) that is context-free, even though the KRAS PC-Map exhibits significant tumor-context specificity (**Figure 5**).

Although PC-Maps allow for the discovery of novel gene proteins in oncogenic signaling, a key limitation is that most gold standards are context-free, thus hampering the ability to generate tissue-specific predictions. The choice of classification methodology is also significant. Naïve Bayes generally underperforms in classification performance and is not feasible when many features are available or correlated features are used. However, they allow direct tracing of the evidence leading to the inference of specific PC-Map interactions. Decision trees are weak learners that can produce irregular patterns by overfitting. Random Forest and other ensemble methods, such as Boosting, correct for this, but bias may ensue, especially if a feature contains a few large values whose effects dominate most trees in a forest. Thus, proteins that are not *bona fide* members of a KRAS PC-Map may be predicted because they are incorrectly learned by the Random Forest classifier due to the use of context-independent PGSSs and/or overfitting. These issues can be mitigated by the use or construction of cancer related PGSSs, normalizing scores within a feature, and investigating different combinations of networks, including experimentally determined ones.

The algorithms presented here produce a single score representing the probability that a protein is a functional member of a specific PC-Map. PrePPI is instrumental for dissecting the layer of physical protein-protein interactions in the KRAS-centric PC-Map whereas ARACNe, VIPER and CINDy provide critical tumor-specificity and discriminate KRAS modulators versus effectors. These individual clues can thus be leveraged to assign proteins to the particular molecular-interaction layers schematically depicted in **Figure 1A**. Specifically, as depicted in **Figure 6** and delineated in **Supplemental Files 4 and 5**, many top scoring KRAS PC-Map proteins can be characterized as upstream modulators (orange), physical binding partners (bold), and downstream effectors (blue), thus producing a highly informative topology. Positive and negative differential activity of candidate effector proteins in the presence of KRAS activating mutations are represented with “+” and “–” symbols, respectively, as determined by VIPER NES values. It is important to note that **Figure 6** shows only the most significant of the KRAS PC-Map interactions to avoid cluttering the image, but there are many similar events uncovered by the analysis. **Supplemental File 5** provides the component feature scores underlying these events. In addition, similarly informative PC-Maps can be constructed for the other oncogenes and oncogenic pathway proteins considered as well as essentially any protein.

An additional critical aspect of PC-Maps represents a fourth functionally important interaction layer (gray lines, **Figure 1A**). Proteins identified as both upstream modulators (CINDy) and downstream effectors (VIPER) of KRAS activity are likely to participate in autoregulatory loops. Indeed, CINDy-inferred upstream modulators can be further assessed as downstream transcriptional targets, for instance, by using the ARACNe algorithm^22^. This can help elucidate the complex autoregulatory circuitry, which is necessary to ensure the stability of cellular phenotypes and may be responsible for complex adaptive behavior, such as in response to pharmacological perturbations.

Two examples of the power of PC-Maps to identify potential autoregulatory networks are described here. HDAC1, a chromatin remodeling enzyme and transcriptional regulator (**Figure 6**, pink oval), is predicted by CINDy as a modulator of two predicted upstream KRAS regulators (JAK2 and ROCK2) and two predicted KRAS effectors (RAPGEF2, RAB8B), denoted by dotted pink lines. Indeed, HDAC1 is overexpressed in lung and other cancers ^76^, and upregulation of HDAC1 affects MYC^27^ and STAT3 protein interactions^77^. Since JAK2-STAT3 signaling regulates many aspects of cancer development and progression^78, 79^ and aberrant MYC activity is induced in KRAS mutant tumors, these functional interactions may pinpoint candidate mechanisms for KRAS-mediated JAK2 activation of STAT3 and direct activation of MYC.

As depicted in **Figure 6**, the dual specificity tyrosine phosphorylated kinase DYRK1B (gray oval) is predicted to aberrantly regulate KRAS. The connection between DYRK1B and KRAS signaling is established^54^ but poorly understood^80–83^. A number of studies have identified DYRK1B as a downstream effector of oncogenic KRAS^80, 81^. However, in support of our prediction, other studies have reported that DYRK1B may function as an upstream regulator of KRAS through its modulation of the mTOR/AKT and MAPK pathways^83^. Oncogenic KRAS promotes hedgehog (HH) signaling ^84^, and HH signaling induces DYRK1B expression, which results in the activation of mTOR/AKT signaling ^83^. Thus, DYRK1B may be subject to complex feedback loops.

As illustrated by the dotted gray lines in **Figure 6**, DYRK1B is predicted to be a modulator of both upstream regulators (IQGAP3) and downstream effectors (e.g. RAP1A and RACGAP1) of KRAS. In support of these predictions, IQGAP3 binds RAS in proliferating cells, and its knockdown decreases RAS activity^85^, while RAP1A antagonistically competes with RAS proteins for binding partners^86^. RACGAP1 activates RAC1, which has been shown to activate the MAPK pathway in a RAS-dependant manner. This suggests that DYRK1B is a partner with KRAS in the regulation of well-established RAS-related pathways, as indicated above. Our predictions, thus, provide testable hypotheses of the possible roles for DYRK1B in mediating these interactions.

Targeting KRAS synthetic lethal (SL) interaction partners is an approach for discovering novel therapeutics in activated KRAS^Mut^ dependent cancers^87^. Predictions of KRAS PC-Map members were highly enriched for KRAS SL interactors, indicating that an appreciable number of KRAS SL partners participate in KRAS signaling (**Figures 2, 3 and 6**). Thus, OncoSig may provide an additional set of pharmacologically accessible targets to target KRAS-mutated tumors. For example, ABL1, validated as a KRAS partner in our assay, is targeted by Imatinib ^88^. Although the role of ABL1 is not well understood in carcinomas such as lung cancer^89^, it may participate in lung cancer metastasis independent of the BCR-ABL1 genomic alteration^90^. ABL1 is activated by RIN1, a RAS interaction partner^91^ with a predicted score of 0.75. Thus, RIN1 may represent a potential alternate target for combination therapy. Among the highest ranked novel predictions (within the top 0.2%, **Supplemental File 4**) are druggable targets discovered in other contexts: 1) LIMK1/Dabrafenib, 2) ITK/Pazopanib, and 3) FYN/Dasatinib, all of which have completed Stage II clinical trials^92^.

However, SL partners depend significantly on tumor type. For example, both Wang et al.^93^ and Barbie et al.^54^ identify CRAF as SL with KRAS^Mut^. Yet, the combination of rigosertib, a RAS mimetic that disrupts CRAF binding to RAS, and gemcitabine, a widely used anti-cancer drug, failed to increase median survival beyond treatment with gemcitabine alone in phase II/III clinical trials for metastatic pancreatic adenocarcinoma KRAS-mutant patients^31^. Indeed, CRAF does not participate in KRAS-driven signaling in pancreatic adenocarcinomas^94^. It is possible that these KRAS-driven tumors acquire resistance to rigosertib by relying on alternative KRAS signaling partners, as depicted in **Figure 6**. A number of previous studies^54, 55, 93, 95, 96^ used high-throughput screens to discover KRAS^mut^ SL interactions, although the overlap of KRAS^Mut^ SL genes reported in these manuscripts is poor ^54, 87^ and do not overlap with the OncoSig_NB_ candidates validated in our assay. However, the enrichment of OncoSig_RF_ predictions in KRAS^Mut^ synthetic lethal proteins identified by other studies (**Figure 4**) and the high validation rate achieved here (**Figure 2**) suggest that many additional bona fide modulators and effectors of KRAS function may be identified even among predictions with lower scores (**Supplemental File 4**), thus further increasing the repertoire of druggable KRAS signaling partners. Our results constitute a valuable resource for guiding high-throughput computational and experimental chemical screens.

OncoSig predicted highly distinct LUAD and COAD-specific KRAS-centric PC-Maps. Although 1,829 high confidence KRAS PC-Map members (S_RF_ ≥ 0.5) were predicted across both contexts, most of them (1,163) of them were uniquely predicted in only one of the two (**Figure 5**) and were thus expected to affect KRAS signaling in context-specific fashion. For example: 1) CABYR (**Figure 5C**, magenta dot), the calcium-binding tyrosine phosphorylation-regulated protein, is overexpressed in lung cancer tumor samples and lung cancer cell-lines^97^, and its knockdown increases sensitivity to drug-induced apoptosis^98^. It is transcriptionally repressed in colon cancer cell lines, and knockdown of its repressor increases its expression ^99^, elevates apoptosis, and suppresses proliferation^100^. 2) While the function of IFITM1 (**Figure 5C**, cyan dot), the interferon-induced transmembrane protein 1, is not known to be associated with KRAS signaling, its locus is deleted in lung cancers^101^, and overexpressed in colorectal cancers where it is associated with poor prognosis^102^. Although the expression levels of these proteins is inherent in the appropriate tumor-specific gene expression profiles, their detection as context-dependent members of the KRAS PC-Map is a distinctive strength of the proposed methodology.

Taken together, the results presented here show that both Naïve Bayes and Random Forest based OncoSig may provide valuable and complementary information to elucidate the complex regulatory machinery that supports the pathophysiologic function of a specific protein or of a set of related proteins in a given cellular context.

## METHODS SUMMARY

Lung adenocarcinoma (LUAD), lung squamous cell carcinoma (LUSC) and colon adenocarcinoma (COAD) gene expression datasets (N_samples_=488, 482 and 434 respectively) were retrieved from The Cancer Genome Atlas (TCGA) and normalized as previously described^24^. We collated 1,813 transcription factors and transcriptional regulators (TFs), 969 transcriptional cofactors (coTFs), and 3,370 signaling proteins (SPs) as described^24^.

### Naive Bayes Classification

To ascertain upstream regulators of KRAS, we inferred the activity of KRAS in LUAD samples using VIPER^24^ and computed two-tailed Normalized Enrichment Score of the KRAS activity. aREA^24^ was used to assess the statistical significance of the co-segregation between nonsynonymous (missense) Single Nucleotide Polymorphisms in other genes and KRAS activity. To identify downstream effectors of aberrant KRAS signaling, we used VIPER to infer the differential activity of TFs, CoTFs, and SPs in KRAS^mut^ samples and closest (based on Spearman correlation) matched KRAS^WT^ samples. A differential gene expression signature ΔEi was computed for each matched KRAS^mut^/KRAS^WT^ pair and the activity change for each TF/CoTF/SP was calculated. Bonferroni-corrected p-values were integrated, using Stouffer’s method producing a p-value for the co-segregation of KRAS^mut^ and the activity of other proteins.

The DeMAND algorithm^25^ was used to discover proteins with dysregulated interactions in KRAS^WT^ versus KRAS^mut^ LUAD samples using a context-specific LUAD molecular interaction network that we previously developed^25, 26^. For each protein, DeMAND predicts which of its interactions are disrupted in KRAS^mut^ versus KRAS^WT^ samples.

MINDy was used to predict post-translational modifications of TFs by SPs, as previously described^27, 62^. A Fisher Exact Test was performed between TFs predicted to be regulated by KRAS and TFs predicted to be regulated by SPs. Each SP was thus assigned a p-value representing the statistical significance of the overlap between the TFs KRAS is predicted to regulate and the TFs other signaling molecules are predicted to regulate.

Predictions of KRAS protein-protein interactions were retrieved from the PrePPI database^28^. Each prediction has an associated Likelihood Ratio (LR) representing the odds above random of the protein-protein interaction occurring.

75 samples with KRAS knockdowns (KDs) in A549 cell lines were retrieved from The Library of Network-Based Cellular Signatures (LINCS) project^29^ (http://www.lincsproject.org/). Averaging over all 75 samples, a single gene expression profile was obtained for each gene.

The NB classifier was trained on a set of 350 proteins annotated as participating in KRAS signaling pathways by the Ingenuity Pathway Analysis. Each clue was split into bins, which were populated by the raw evidence values such that an equal number of members of the positive gold standard set (PGSS) was distributed across bins as possible. Training was performed using two-fold cross validation with holdout, which creates an independent training and testing set and produces a final LR for every protein that is parameterized on the set to which it does not belong.

### Random Forest Classification

The ARACNe, VIPER and CINDY algorithms were applied to LUAD, LUSC and COAD gene expression profiles with the same parameters and p-value thresholds as previously described^24, 62^. Protein-protein interactions from the PrePPI database were retrieved. The LR_GO_ and LR_Exp_ components were removed from LR_PrePPI_, and interactions with modified LR_PrePPI_ scores ≥ 600 were used.

The PGSSs for the 10 pathways used in RF classification were compiled as the union of the KEGG, Biocarta, and Reactome databases from the MSigDB C2 category^65^ and further pathway members from the KEGG website^5^ KRAS synthetic lethality and drug-dependency data were compiled from Barbie et al.^54^, Corcoran et al.^70^, Hayes et al.^71^, Astsaturov et al.^73^ and GO:0000302: “Response to Reactive Oxygen Species”^6^. Protein domain data was obtained from PFAM version 31^103^. CD-HIT^104^ was used to generate a Non-Redundant KRAS PGSS. The KRAS PGSS was clustered at an 80% sequence identity threshold and, for each cluster, the representative with the longest sequence was selected as per CD-HIT protocol.

The features derived from the networks were as follows: Mutual information for ARACNe, number of statistically significant triplets for CINDy, negative log p-value for VIPER, and LR for PrePPI. We coded each feature symmetrically, so that interactions between protein A and protein B were input into the matrix twice, once in the feature vector for A and once in the feature vector for B; all other elements in the in the Random Forest feature matrix were set to zero. For each of the 10 oncogene-centric interactomes, proteins that are part of the PGSS were assigned a “1” within the PGSS vector, while all other proteins were assigned a “0” to represent membership in the NGSS. Training and testing of the Random Forest classifier then proceeded using each pathway’s PGSS and NGSS using Monte Carlo cross-validation^105^, creating 50 forests each with 50 trees . We performed 100 OncoSig runs each with the LUAD, COAD and LUSC KRAS PGSS networks and distribution of Pearson correlation coefficients were estimated by calculating all pairwise Pearson correlation coefficients.

### Tandem Affinity Purification

5 mL packed cell volume of RPE-hTERT cells expressing LAP-tagged proteins were resuspended with 20 mL of LAP-resuspension buffer, lysed, and then incubated on ice for 10 min. The lysate was first centrifuged at 14,000 rpm (27,000 g) at 4°C for 10 min, and the resulting supernatant was centrifuged at 43,000 rpm (100,000 g) for 1 hr at 4°C to further clarify the lysate. High speed supernatant was mixed with 500 µL of GFP-coupled beads^30^ and rotated for 1 hr at 4°C to capture GFP-tagged proteins, and washed five times with 1 mL LAP200N. After re-suspending the beads with 1 mL LAP200N buffer lacking DTT and protease inhibitors, the GFP-tag was cleaved by adding 5 µg of TEV protease and rotating tubes at 4°C overnight. TEV-eluted supernatant was added to 100 µL of S-protein agarose to capture S-tagged protein. After washing three times with LAP200N buffer lacking DTT and twice with LAP100 buffer, purified protein complexes were eluted with 50 µL of 2X LDS buffer and boiled at 95°C for 3 min. Samples were then run on Bolt® Bis-Tris Plus Gels in Bolt® MES SDS Running Buffer. Gels were fixed in 100 mL of fixing solution at room temperature, and stained with Colloidal Blue Staining Kit. After the buffer was replaced with Optima™ water, the bands were cut into eight pieces, followed by washing twice with 500 µL of 50% acetonitrile in Optima™ water. The gel slices were then reduced and alkylated followed by destaining and in-gel digestion using 125 ng Trypsin/LysC as previously described^106^. Tryptic peptides were extracted from the gel bands and dried in a speed vac. Prior to LC-MS, each sample was reconstituted in 0.1% formic acid, 2% acetonitrile, and water. NanoAcquity (Waters) LC instrument was set at a flow rate of either 300 nL/min or 450 nL/min where mobile phase A was 0.2% formic acid in water and mobile phase B was 0.2% formic acid in acetonitrile. The analytical column was in-house pulled and packed using C18 Reprosil Pur 2.4 uM where the I.D. was 100 uM and the column length was 20-25 cm. Peptide pools were directly injected onto the analytical column in which linear gradients (4-40% B) were of either 80 or 120 min eluting peptides into the mass spectrometer. MS/MS was acquired using CID with a collisional energy of 32-35. In a typical analysis, RAW files were processed using Byonic (Protein Metrics) using 12 ppm mass accuracy limits for precursors and 0.4 Da mass accuracy limits for MS/MS spectra. MS/MS data was compared to an NCBI Genbank FASTA database containing all human proteomic isoforms with the exception of the tandem affinity bait construct sequence and common contaminant proteins. Spectral counts were assumed to have undergone fully specific proteolysis and allowing up to two missed cleavages per peptide.

### Primary Tumor Propagating Cell Culture and Screening Methodology

Primary lung tumor cells from KRAS^G12D/+^; p53^fl/fl^ mice were cultured in Matrigel as described previously^52^. Prior to seeding, primary cells were infected with a pool of 100-150 lentiviral pLKO shRNAs composed of 3-5 shRNAs per gene at a Multiplicity of Infection <0.5 to ensure single shRNA integration and selected with 1ug/ml puromycin 24 hours after seeding. We screened the top 22 predicted genes from the NB classifier in two pools. Pools also included other candidate vulnerabilities identified by literature review and other methods. 25 pools consisting of 2,286 shRNAs targeting 515 genes not anticipated to be involved in KRAS-regulated signaling were used as a background comparison. After 7 days of spheroid growth, spheroids were dissociated with trypsin into single cells, and half of the 3D culture was re-seeded. The remaining half of each sample was retained for gDNA isolation (T0) until secondary spheroids fully formed 7 days later (T1). The integrated pLKO shRNA was PCR amplified using ExTaq (Clontech), barcoded, multiplexed, and sequenced on an Illumina GAIIx (primer sequences available on request). Sequencing reads were processed into count files in R (v. 3.1.1) using the edgeR package (v. 3.6.8) and analyzed using generalized linear models with edgeR using a time-course design to compare the initial (T0) and final (T1) timepoints and perform a likelihood ratio test^107^.

To calculate the statistical significance of the fold change in growth induced by each individual shRNA, we fit a density plot of all the background screens. For each shRNA, we integrated from the minimum log_2_FC of the entire BPS to the log_2_FC observed for that shRNA, producing a one-tailed p-value for the observed log_2_FC. We used Fisher’s method to integrate the p-values of all shRNAs that mapped to the same protein.

### Gene Enrichment Analysis

Gene enrichment analysis was done by extracting all GO Biological Process terms from the PANTHER database^108^, and, for each GO term, testing for overrepresentation between the Naïve Bayes candidates with an integrated p-value<=.05 and 468 members of BPS that were represented in the human Uniprot proteome.

### Multiple Hypothesis Correction

All p-values reported for all analyses (except where noted otherwise) were corrected using the Benjamini & Hochberg False Discovery Rate^58^.

### Data and Code Availability

The Online Methods and accompanying codebook contain a further description of the analyses performed. Code for running Oncosig-NB, Oncosig-RF and perform statistical analysis of the pooled shRNA results are provided in the accompanying codebook.

## Acknowledgements

This work was supported by the NCI Outstanding Investigator Award R35CA197745 to AC; the NCI Cancer Target Discovery and Development Program U01CA168426 to A.C.; the NCI Research Centers for Cancer Systems Biology Consortium U54CA209997 to A.C. and B.H.; NIGMS grant R01GM30518 to B.H.; NCI grant R01CA129562 to E.A.S.C.; Innovative Research Grant from Stand up to Cancer to E.A.S.C.; NIH High-end Instrumentation Program grant S10OD012351 to A.C.; the NIH Shared Instrumentation Program grant S10OD021764 to A.C.

J.B. was supported in part by the Ruth L. Kirschstein National Research Service Award Institutional Research Training Grant T32GM082797. D.R.S. was supported by the Ruth L. Kirschstein National Research Service Award Institutional Research Training Grant T32CA09302.

## Author Contributions

J.B., A.L. and F.M.G. performed computational analysis. J.B. performed machine learning. D.S. and P.K.J. performed, respectively, the knockdown and AP/MS experiments. J.B., D.S., E.A.S.C., A.C. and B.H. analyzed knockdown experiments. D.M., B.H., A.C., and E.A.S.C. designed research. D.M. B.H and A.C designed computational and experimental work and analyzed data. D.M., D.S. J.B., A.C., and B.H assembled the data and wrote the paper.

## REFERENCES

1. Papin, J. A., Hunter, T., Palsson, B. O. & Subramaniam, S. Reconstruction of cellular signalling networks and analysis of their properties. Nat Rev Mol Cell Biol 6, 99–111, doi:10.1038/nrm1570 (2005).

2. Springer, M. S., Goy, M. F. & Adler, J. Protein methylation in behavioural control mechanisms and in signal transduction. Nature 280, 279–284 (1979).

3. Wiley, H. S., Shvartsman, S. Y. & Lauffenburger, D. A. Computational modeling of the EGF-receptor system: a paradigm for systems biology. Trends in cell biology 13, 43–50 (2003).

4. Califano, A., Butte, A. J., Friend, S., Ideker, T. & Schadt, E. Leveraging models of cell regulation and GWAS data in integrative network-based association studies. Nat Genet 44, 841–847, doi:10.1038/ng.2355 (2012).

5. Kanehisa, M., Sato, Y., Kawashima, M., Furumichi, M. & Tanabe, M. KEGG as a reference resource for gene and protein annotation. Nucleic Acids Res 44, D457–462, doi:10.1093/nar/gkv1070 (2016).

6. Ashburner, M. et al. Gene ontology: tool for the unification of biology. The Gene Ontology Consortium. Nat Genet 25, 25–29, doi:10.1038/75556 (2000).

7. Nishimura, D. BioCarta. Biotech Software & Internet Report 2, 117–120 (2001).

8. Croft, D. et al. The Reactome pathway knowledgebase. Nucleic Acids Res 42, D472–477, doi:10.1093/nar/gkt1102 (2014).

9. Paz, A. et al. SPIKE: a database of highly curated human signaling pathways. Nucleic Acids Res 39, D793–799, doi:10.1093/nar/gkq1167 (2011).

10. Cerami, E. G. et al. Pathway Commons, a web resource for biological pathway data. Nucleic Acids Res 39, D685–690, doi:10.1093/nar/gkq1039 (2011).

11. Kramer, A., Green, J., Pollard, J., Jr. & Tugendreich, S. Causal analysis approaches in Ingenuity Pathway Analysis. Bioinformatics 30, 523–530, doi:10.1093/bioinformatics/btt703 (2014).

12. Prahallad, A. et al. Unresponsiveness of colon cancer to BRAF(V600E) inhibition through feedback activation of EGFR. Nature 483, 100–103, doi:10.1038/nature10868 (2012).

13. Rozengurt, E., Soares, H. P. & Sinnet-Smith, J. Suppression of feedback loops mediated by PI3K/mTOR induces multiple overactivation of compensatory pathways: an unintended consequence leading to drug resistance. Molecular cancer therapeutics 13, 2477–2488, doi:10.1158/1535-7163.mct-14-0330 (2014).

14. Haarberg, H. E. & Smalley, K. S. Resistance to Raf inhibition in cancer. Drug Discov Today Technol 11, 27–32, doi:10.1016/j.ddtec.2013.12.004 (2014).

15. Karnoub, A. E. & Weinberg, R. A. Ras oncogenes: split personalities. Nature reviews 9, 517–531, doi:10.1038/nrm2438 (2008).

16. Villaruz, L. C. et al. The prognostic and predictive value of KRAS oncogene substitutions in lung adenocarcinoma. Cancer 119, 2268–2274, doi:10.1002/cncr.28039 (2013).

17. Bhattacharya, S., Socinski, M. A. & Burns, T. F. KRAS mutant lung cancer: progress thus far on an elusive therapeutic target. Clin Transl Med 4, 35, doi:10.1186/s40169-015-0075-0 (2015).

18. Haigis, K. M. et al. Differential effects of oncogenic K-Ras and N-Ras on proliferation, differentiation and tumor progression in the colon. Nat Genet 40, 600–608, doi:10.1038/ng.115 (2008).

19. Jansen, R. et al. A Bayesian networks approach for predicting protein-protein interactions from genomic data. Science 302, 449–453, doi:10.1126/science.1087361302/5644/449 [pii] (2003).

20. Breiman, L. Random forests. Mach Learn 45, 5–32, doi:Doi 10.1023/A:1010933404324 (2001).

21. Network, C. G. A. R. Comprehensive molecular profiling of lung adenocarcinoma. Nature 511, 543–550, doi:10.1038/nature13385 (2014).

22. Margolin, A. A. et al. ARACNE: an algorithm for the reconstruction of gene regulatory networks in a mammalian cellular context. BMC Bioinformatics 7 Suppl 1, S7, doi:10.1186/1471-2105-7-s1-s7 (2006).

23. Schadt, E. E. et al. An integrative genomics approach to infer causal associations between gene expression and disease. Nat Genet 37, 710–717, doi:10.1038/ng1589 (2005).

24. Alvarez, M. J. et al. Functional characterization of somatic mutations in cancer using network-based inference of protein activity. Nat Genet 48, 838–847, doi:10.1038/ng.3593 (2016).

25. Woo, J. H. et al. Elucidating Compound Mechanism of Action by Network Perturbation Analysis. Cell 162, 441–451, doi:10.1016/j.cell.2015.05.056 (2015).

26. Lefebvre, C. et al. A human B-cell interactome identifies MYB and FOXM1 as master regulators of proliferation in germinal centers. Mol Syst Biol 6, 377, doi:10.1038/msb.2010.31 (2010).

27. Wang, K. et al. Genome-wide identification of post-translational modulators of transcription factor activity in human B cells. Nat Biotechnol 27, 829–839, doi:nbt.1563 [pii] 10.1038/nbt.1563 (2009).

28. Zhang, Q. C. et al. Structure-based prediction of protein-protein interactions on a genome-wide scale. Nature 490, 556–560, doi:10.1038/nature11503 (2012).

29. Duan, Q. et al. LINCS Canvas Browser: interactive web app to query, browse and interrogate LINCS L1000 gene expression signatures. Nucleic Acids Res 42, W449–460, doi:10.1093/nar/gku476 (2014).

30. Torres, J. Z., Miller, J. J. & Jackson, P. K. High-throughput generation of tagged stable cell lines for proteomic analysis. Proteomics 9, 2888–2891, doi:10.1002/pmic.200800873 (2009).

31. (QIAGEN’s Ingenuity® Pathway Analysis, IPA®, QIAGEN Redwood City, www.qiagen.com/ingenuity).

32. Scheffzek, K. et al. The Ras-RasGAP complex: structural basis for GTPase activation and its loss in oncogenic Ras mutants. Science 277, 333–338 (1997).

33. Villalonga, P. et al. Calmodulin binds to K-Ras, but not to H- or N-Ras, and modulates its downstream signaling. Mol Cell Biol 21, 7345–7354, doi:10.1128/mcb.21.21.7345-7354.2001 (2001).

34. Kupzig, S. et al. GAP1 family members constitute bifunctional Ras and Rap GTPase-activating proteins. The Journal of biological chemistry 281, 9891–9900, doi:10.1074/jbc.M512802200 (2006).

35. Beaudoin, G. M., 3rd et al. Afadin, a Ras/Rap effector that controls cadherin function, promotes spine and excitatory synapse density in the hippocampus. The Journal of neuroscience : the official journal of the Society for Neuroscience 32, 99–110, doi:10.1523/jneurosci.4565-11.2012 (2012).

36. Duran, A. et al. The signaling adaptor p62 is an important NF-kappaB mediator in tumorigenesis. Cancer Cell 13, 343–354, doi:10.1016/j.ccr.2008.02.001 (2008).

37. Fetics, S. K. et al. Allosteric effects of the oncogenic RasQ61L mutant on Raf-RBD. Structure 23, 505–516, doi:10.1016/j.str.2014.12.017 (2015).

38. Kidd, A. R., 3rd et al. Ras-related small GTPases RalA and RalB regulate cellular survival after ionizing radiation. Int J Radiat Oncol Biol Phys 78, 205–212, doi:10.1016/j.ijrobp.2010.03.023 (2010).

39. Kitayama, H., Sugimoto, Y., Matsuzaki, T., Ikawa, Y. & Noda, M. A ras-related gene with transformation suppressor activity. Cell 56, 77–84 (1989).

40. Lambert, J. M. et al. Tiam1 mediates Ras activation of Rac by a PI(3)K-independent mechanism. Nature cell biology 4, 621–625, doi:10.1038/ncb833 (2002).

41. Qiu, R. G., Abo, A., McCormick, F. & Symons, M. Cdc42 regulates anchorage-independent growth and is necessary for Ras transformation. Mol Cell Biol 17, 3449–3458 (1997).

42. Rusanescu, G., Gotoh, T., Tian, X. & Feig, L. A. Regulation of Ras signaling specificity by protein kinase C. Mol Cell Biol 21, 2650–2658, doi:10.1128/mcb.21.8.2650-2658.2001 (2001).

43. Stewart, S. et al. Kinase suppressor of Ras forms a multiprotein signaling complex and modulates MEK localization. Mol Cell Biol 19, 5523–5534 (1999).

44. Tall, G. G., Barbieri, M. A., Stahl, P. D. & Horazdovsky, B. F. Ras-activated endocytosis is mediated by the Rab5 guanine nucleotide exchange activity of RIN1. Developmental cell 1, 73–82 (2001).

45. Zhou, F. et al. The mechanism and function of mitogen-activated protein kinase activation by ARF1. Cellular signalling 27, 2035–2044, doi:10.1016/j.cellsig.2015.06.007 (2015).

46. Chatr-Aryamontri, A. et al. The BioGRID interaction database: 2013 update. Nucleic Acids Res 41, D816–823, doi:10.1093/nar/gks1158 (2013).

47. Matsunaga, H., Kubota, K., Inoue, T., Isono, F. & Ando, O. IQGAP1 selectively interacts with K-Ras but not with H-Ras and modulates K-Ras function. Biochem Biophys Res Commun 444, 360–364, doi:10.1016/j.bbrc.2014.01.041 (2014).

48. Fatehullah, A., Tan, S. H. & Barker, N. Organoids as an in vitro model of human development and disease. Nat Cell Biol 18, 246–254, doi:10.1038/ncb3312 (2016).

49. Pampaloni, F., Reynaud, E. G. & Stelzer, E. H. The third dimension bridges the gap between cell culture and live tissue. Nat Rev Mol Cell Biol 8, 839–845, doi:10.1038/nrm2236 (2007).

50. Wang, F. et al. Reciprocal interactions between beta1-integrin and epidermal growth factor receptor in three-dimensional basement membrane breast cultures: a different perspective in epithelial biology. Proc Natl Acad Sci U S A 95, 14821–14826 (1998).

51. Weaver, V. M. et al. beta4 integrin-dependent formation of polarized three-dimensional architecture confers resistance to apoptosis in normal and malignant mammary epithelium. Cancer Cell 2, 205–216 (2002).

52. Zheng, Y. et al. A rare population of CD24(+)ITGB4(+)Notch(hi) cells drives tumor propagation in NSCLC and requires Notch3 for self-renewal. Cancer Cell 24, 59–74, doi:10.1016/j.ccr.2013.05.021 (2013).

53. Papke, B. & Der, C. J. Drugging RAS: Know the enemy. Science 355, 1158–1163, doi:10.1126/science.aam7622 (2017).

54. Barbie, D. A. et al. Systematic RNA interference reveals that oncogenic KRAS-driven cancers require TBK1. Nature 462, 108–112, doi:10.1038/nature08460 (2009).

55. Kim, J. et al. XPO1-dependent nuclear export is a druggable vulnerability in KRAS-mutant lung cancer. Nature 538, 114–117, doi:10.1038/nature19771 (2016).

56. Shrestha, G. et al. The value of genomics in dissecting the RAS-network and in guiding therapeutics for RAS-driven cancers. Seminars in cell & developmental biology 58, 108–117, doi:10.1016/j.semcdb.2016.06.012 (2016).

57. Fisher, R. A. Statistical methods for research workers. 4th edn, (Oliver and Boyd, 1932).

58. Benjamini, Y. & Hochberg, Y. Controlling the False Discovery Rate - a Practical and Powerful Approach to Multiple Testing. Journal of the Royal Statistical Society Series B-Methodological 57, 289–300 (1995).

59. Caruana, R. & Niculescu-Mizil, A. An Empirical Comparison of Supervised Learning Algorithms. Proceedings of the 23rd International Conference on Machine Learning 161–168 (2006).

60. Pan, D. The hippo signaling pathway in development and cancer. Dev Cell 19, 491–505, doi:10.1016/j.devcel.2010.09.011 (2010).

61. Polakis, P. Wnt signaling in cancer. Cold Spring Harb Perspect Biol 4, doi:10.1101/cshperspect.a008052 (2012).

62. Giorgi, F. M. et al. Inferring protein modulation from gene expression data using conditional mutual information. PLoS One 9, e109569, doi:10.1371/journal.pone.0109569 (2014).

63. Garzon, J. I. et al. A computational interactome and functional annotation for the human proteome. Elife 5, doi:10.7554/eLife.18715 (2016).

64. Forbes, S. A. et al. COSMIC: exploring the world’s knowledge of somatic mutations in human cancer. Nucleic Acids Res 43, D805–811, doi:10.1093/nar/gku1075 (2015).

65. Liberzon, A. et al. The Molecular Signatures Database (MSigDB) hallmark gene set collection. Cell Syst 1, 417–425, doi:10.1016/j.cels.2015.12.004 (2015).

66. Subramanian, A. et al. Gene set enrichment analysis: a knowledge-based approach for interpreting genome-wide expression profiles. Proc Natl Acad Sci U S A 102, 15545–15550, doi:10.1073/pnas.0506580102 (2005).

67. Li, W. & Godzik, A. Cd-hit: a fast program for clustering and comparing large sets of protein or nucleotide sequences. Bioinformatics 22, 1658–1659, doi:10.1093/bioinformatics/btl158 (2006).

68. Franceschini, A. et al. STRING v9.1: protein-protein interaction networks, with increased coverage and integration. Nucleic Acids Res 41, D808–815, doi:10.1093/nar/gks1094 (2013).

69. Lee, I., Blom, U. M., Wang, P. I., Shim, J. E. & Marcotte, E. M. Prioritizing candidate disease genes by network-based boosting of genome-wide association data. Genome Res 21, 1109–1121, doi:10.1101/gr.118992.110 (2011).

70. Corcoran, R. B. et al. Synthetic lethal interaction of combined BCL-XL and MEK inhibition promotes tumor regressions in KRAS mutant cancer models. Cancer Cell 23, 121–128, doi:10.1016/j.ccr.2012.11.007 (2013).

71. Hayes, T. K. et al. Long-Term ERK Inhibition in KRAS-Mutant Pancreatic Cancer Is Associated with MYC Degradation and Senescence-like Growth Suppression. Cancer Cell 29, 75–89, doi:10.1016/j.ccell.2015.11.011 (2016).

72. Shaw, A. T. et al. Selective killing of K-ras mutant cancer cells by small molecule inducers of oxidative stress. Proc Natl Acad Sci U S A 108, 8773–8778, doi:10.1073/pnas.1105941108 (2011).

73. Astsaturov, I. et al. Synthetic lethal screen of an EGFR-centered network to improve targeted therapies. Sci Signal 3, ra67, doi:10.1126/scisignal.2001083 (2010).

74. Lannon, C. L. & Sorensen, P. H. ETV6-NTRK3: a chimeric protein tyrosine kinase with transformation activity in multiple cell lineages. Semin Cancer Biol 15, 215–223, doi:10.1016/j.semcancer.2005.01.003 (2005).

75. Stengel, K. R. & Zheng, Y. Essential role of Cdc42 in Ras-induced transformation revealed by gene targeting. PLoS One 7, e37317, doi:10.1371/journal.pone.0037317 (2012).

76. Barneda-Zahonero, B. & Parra, M. Histone deacetylases and cancer. Mol Oncol 6, 579–589, doi:10.1016/j.molonc.2012.07.003 (2012).

77. Singh, B. N. et al. Nonhistone protein acetylation as cancer therapy targets. Expert Rev Anticancer Ther 10, 935–954, doi:10.1586/era.10.62 (2010).

78. Teng, Y., Ross, J. L. & Cowell, J. K. The involvement of JAK-STAT3 in cell motility, invasion, and metastasis. JAKSTAT 3, e28086, doi:10.4161/jkst.28086 (2014).

79. Yu, H., Lee, H., Herrmann, A., Buettner, R. & Jove, R. Revisiting STAT3 signalling in cancer: new and unexpected biological functions. Nat Rev Cancer 14, 736–746, doi:10.1038/nrc3818 (2014).

80. Jin, K., Park, S., Ewton, D. Z. & Friedman, E. The survival kinase Mirk/Dyrk1B is a downstream effector of oncogenic K-ras in pancreatic cancer. Cancer Res 67, 7247–7255, doi:10.1158/0008-5472.CAN-06-4099 (2007).

81. Lauth, M. et al. DYRK1B-dependent autocrine-to-paracrine shift of Hedgehog signaling by mutant RAS. Nature structural & molecular biology 17, 718–725, doi:10.1038/nsmb.1833 (2010).

82. Gruber, W. et al. DYRK1B as therapeutic target in Hedgehog/GLI-dependent cancer cells with Smoothened inhibitor resistance. Oncotarget 7, 7134–7148, doi:10.18632/oncotarget.6910 (2016).

83. Singh, R., Dhanyamraju, P. K. & Lauth, M. DYRK1B blocks canonical and promotes non-canonical Hedgehog signaling through activation of the mTOR/AKT pathway. Oncotarget 8, 833–845, doi:10.18632/oncotarget.13662 (2017).

84. Ji, Z., Mei, F. C., Xie, J. & Cheng, X. Oncogenic KRAS activates hedgehog signaling pathway in pancreatic cancer cells. The Journal of biological chemistry 282, 14048–14055, doi:10.1074/jbc.M611089200 (2007).

85. Nojima, H. et al. IQGAP3 regulates cell proliferation through the Ras/ERK signalling cascade. Nat Cell Biol 10, 971–978, doi:10.1038/ncb1757 (2008).

86. Bos, J. L., Franke, B., M’Rabet, L., Reedquist, K. & Zwartkruis, F. In search of a function for the Ras-like GTPase Rap1. FEBS letters 410, 59–62 (1997).

87. Downward, J. RAS Synthetic Lethal Screens Revisited: Still Seeking the Elusive Prize? Clinical cancer research : an official journal of the American Association for Cancer Research 21, 1802–1809, doi:10.1158/1078-0432.ccr-14-2180 (2015).

88. Nagar, B. c-Abl tyrosine kinase and inhibition by the cancer drug imatinib (Gleevec/STI-571). J Nutr 137, 1518S–1523S; discussion 1548S (2007).

89. Greuber, E. K., Smith-Pearson, P., Wang, J. & Pendergast, A. M. Role of ABL family kinases in cancer: from leukaemia to solid tumours. Nat Rev Cancer 13, 559–571, doi:10.1038/nrc3563 (2013).

90. Gu, J. J. et al. Inactivation of ABL kinases suppresses non-small cell lung cancer metastasis. JCI Insight 1, e89647, doi:10.1172/jci.insight.89647 (2016).

91. Hu, H., Bliss, J. M., Wang, Y. & Colicelli, J. RIN1 is an ABL tyrosine kinase activator and a regulator of epithelial-cell adhesion and migration. Current biology : CB 15, 815–823, doi:10.1016/j.cub.2005.03.049 (2005).

92. Koscielny, G. et al. Open Targets: a platform for therapeutic target identification and validation. Nucleic Acids Res 45, D985–D994, doi:10.1093/nar/gkw1055 (2017).

93. Wang, T. et al. Gene Essentiality Profiling Reveals Gene Networks and Synthetic Lethal Interactions with Oncogenic Ras. Cell 168, 890–903 e815, doi:10.1016/j.cell.2017.01.013 (2017).

94. Eser, S., Schnieke, A., Schneider, G. & Saur, D. Oncogenic KRAS signalling in pancreatic cancer. Br J Cancer 111, 817–822, doi:10.1038/bjc.2014.215 (2014).

95. Luo, J. et al. A genome-wide RNAi screen identifies multiple synthetic lethal interactions with the Ras oncogene. Cell 137, 835–848, doi:10.1016/j.cell.2009.05.006 (2009).

96. McDonald, E. R., 3rd et al. Project DRIVE: A Compendium of Cancer Dependencies and Synthetic Lethal Relationships Uncovered by Large-Scale, Deep RNAi Screening. Cell 170, 577–592 e510, doi:10.1016/j.cell.2017.07.005 (2017).

97. Tseng, Y. T. et al. Expression of the sperm fibrous sheath protein CABYR in human cancers and identification of alpha-enolase as an interacting partner of CABYR-a. Oncol Rep 25, 1169–1175, doi:10.3892/or.2011.1165 (2011).

98. Qian, Z. et al. Knockdown of CABYR-a/b increases chemosensitivity of human non-small cell lung cancer cells through inactivation of Akt. Mol Cancer Res 12, 335–347, doi:10.1158/1541-7786.mcr-13-0391 (2014).

99. Chen, J. et al. Identification of downstream metastasis-associated target genes regulated by LSD1 in colon cancer cells. Oncotarget 8, 19609–19630, doi:10.18632/oncotarget.14778 (2017).

100. Ding, J. et al. LSD1-mediated epigenetic modification contributes to proliferation and metastasis of colon cancer. Br J Cancer 109, 994–1003, doi:10.1038/bjc.2013.364 (2013).

101. Catchpoole, D. et al. Epigenetic modification and uniparental inheritance of H19 in Beckwith-Wiedemann syndrome. J Med Genet 34, 353–359 (1997).

102. He, J., Li, J., Feng, W., Chen, L. & Yang, K. Prognostic significance of INF-induced transmembrane protein 1 in colorectal cancer. Int J Clin Exp Pathol 8, 16007–16013 (2015).

## METHODS SUMMARY REFERENCES

103. Finn, R. D. et al. The Pfam protein families database: towards a more sustainable future. Nucleic Acids Res 44, D279–285, doi:10.1093/nar/gkv1344 (2016).

104. Li, W. Z. & Godzik, A. Cd-hit: a fast program for clustering and comparing large sets of protein or nucleotide sequences. Bioinformatics 22, 1658–1659, doi:10.1093/bioinformatics/btl158 (2006).

105. Shao, J. Linear-Model Selection by Cross-Validation. Journal of the American Statistical Association 88, 486–494, doi:Doi 10.2307/2290328 (1993).

106. Shevchenko, A., Tomas, H., Havlis, J., Olsen, J. V. & Mann, M. In-gel digestion for mass spectrometric characterization of proteins and proteomes. Nat Protoc 1, 2856–2860, doi:10.1038/nprot.2006.468 (2006).

107. Dai, Z. et al. edgeR: a versatile tool for the analysis of shRNA-seq and CRISPR-Cas9 genetic screens. F1000Res 3, 95, doi:10.12688/f1000research.3928.2 (2014).

108. Mi, H., Poudel, S., Muruganujan, A., Casagrande, J. T. & Thomas, P. D. PANTHER version 10: expanded protein families and functions, and analysis tools. Nucleic Acids Res 44, D336–342, doi:10.1093/nar/gkv1194 (2016).

