## Supplementary Materials for "Systematic Elucidation and Validation of OncoProtein-Centric Molecular Interaction Maps"

### ONLINE METHODS:

Lung adenocarcinoma (LUAD), lung squamous cell carcinoma (LUSC) and colon adenocarcinoma (COAD) gene expression datasets ($N_{samples}$=488, 482 and 434 respectively) were retrieved from The Cancer Genome Atlas (TCGA) and normalized as previously described[24]. For LUAD, 326 had $KRAS^{WT}$, 134 had $KRAS^{mut}$, and 28 had no information on KRAS mutation status and were excluded. Guided by annotation from the Gene Ontology Consortium (GO)[6], we retrieved 1,813 transcription factors and transcriptional regulators (TFs), 969 transcriptional cofactors (coTFs), and 3,370 signaling proteins (SPs).

#### Naive Bayes Classification

#### VIPER

To ascertain upstream regulators of KRAS, we inferred the activity of KRAS in LUAD samples using VIPER[24]. ARACNe [22], with default parameters (p-value threshold p = $10^{-8}$), was used to infer regulons for KRAS and other proteins from LUAD gene expression data. VIPER computed the two-tailed Normalized Enrichment Score (NES) of the KRAS regulon in genes differentially expressed in each sample, compared to the average expression level across all other samples. Thus, positive and negative NES values indicate an increase or decrease in KRAS activity compared to average activity. We then checked whether samples with high VIPER-inferred activity of KRAS co-segregated nonsynonymous (missense) Single Nucleotide Polymorphisms (nsSNPs) in other genes. aREA[24] was used to assess the statistical significance of the co-segregation to obtain an FDR-corrected p-value of the co-segregation between the mutation of other genes and KRAS activity.

To identify downstream effectors of aberrant KRAS signaling, we selected proteins with significant VIPER-inferred activity changes in those LUAD samples with a known KRAS activating mutation. Using ARACNe-inferred regulons, VIPER inferred the differential activity of TFs, CoTFs, and SPs in $KRAS^{mut}$ samples and closest matched $KRAS^{WT}$ samples, where sample proximity was determined by the Spearman correlation between gene expression profiles. $KRAS^{mut}$ samples with inferred KRAS activity below the dataset median were excluded (along with their matched $KRAS^{WT}$ samples), as KRAS tumorigenic signals in these samples may have been attenuated by other events[109]. If $KRAS^{WT}$ samples contained mutations in known RAS pathway genes (i.e., NF1, SOS1, NRAS, HRAS, RAF1, ARAF, BRAF, MAP2K1 and MAPK1) or had inferred KRAS activity above the median, they and their matched $KRAS^{mut}$ samples were excluded. This resulted in a subset of 221 samples. A differential gene expression signature ΔEi was computed for each matched $KRAS^{mut}$/$KRAS^{WT}$ pair. Finally, the relative activity change of each TF, CoTF, and SP was inferred by computing the enrichment of its ARACNe-inferred regulon in genes differentially expressed in ΔEi. Bonferroni-corrected p-values were integrated, using Stouffer's method, to identify global effectors genes. This produced a p-value for the co-segregation of $KRAS^{mut}$ and the activity of other proteins.

**DeMAND**

We used the DeMAND algorithm[25] to discover proteins with dysregulated interactions in $KRAS^{WT}$ versus $KRAS^{mut}$ LUAD samples. We used a context-specific LUAD molecular interaction network that we previously developed which incorporates protein-protein and protein-DNA regulatory networks[25,26]. For each protein, DeMAND predicts which of its interactions are disrupted in $KRAS^{mut}$ versus $KRAS^{WT}$ samples. Briefly, the Kullback-Leibler divergence is calculated for each edge in the network to assess whether it is disrupted in KRAS$^{mut}$ vs. KRAS$^{WT}$ samples. The p-values of a protein's disrupted edges are integrated using Fisher's method and corrected using Brown's method. This produces a final p-value for every protein which represents the dysregulation of its edges between KRAS$^{WT}$ and KRAS$^{mut}$ samples. To avoid biasing DeMAND predictions to previously predicted or known KRAS functional partners, these edges were removed prior to running the algorithm.

**MINDy**

MINDy was used to predict post-translational modifications of TFs by SPs, as previously described[27,62], with a p-value threshold of ≤ 0.005. A Fisher Exact Test was performed between TFs predicted to be regulated by KRAS and TFs predicted to be regulated by SPs. Each SP was thus assigned a p-value representing the statistical significance of the overlap between the TFs KRAS is predicted to regulate and the TFs other signaling molecules are predicted to regulate.

**PrePPI**

Predictions of KRAS protein-protein interactions were retrieved from the PrePPI database[28]. Each prediction has an associated Likelihood Ratio (LR) representing the odds above random of the protein-protein interaction occurring.

**LINCS**

75 samples with KRAS knockdowns (KDs) in A549 cell lines were retrieved from The Library of Network-Based Cellular Signatures (LINCS) project[29] (http://www.lincsproject.org/). The KRAS KDs had been performed with three different hairpins and the response in gene expression was measured after 96 hours, which was z-transformed across experiments. We calculated the change in gene expression in

response to KRAS KD. Averaging over all 75 samples, a single gene expression profile was obtained for each gene, and the log fold change of expression from these average profiles was calculated.

**Parameterization of the Naïve Bayes Classifier**

We implemented a Naïve Bayes (NB) classifier to predict novel members of KRAS regulated pathways. The NB classifier was trained on a set of 350 proteins annotated as participating in KRAS signaling pathways by the Ingenuity Pathway Analysis.

Value types for each clue are LRs for PrePPI, p-values for MINDy, VIPER and DeMAND, log fold change of gene expression in response to KRAS knockdown for LINCS, and peptide count for AP-MS. Each clue was split into bins, which were populated by the raw evidence value such that an approximately equal number of members of the positive gold standard set (PGSS) were distributed across bins. In cases of highly skewed distributions, bin size was fit further to prevent dilution of signal. If a clue did not have a corresponding value for a particular protein, it was assigned to an "NA" bin. $LR_{Bin}$ represents the increased odds that a protein assigned to a particular bin is involved in KRAS signaling as compared to random. Specifically, LR for each bin is given by:

$$LR_{Bin} = (P(PGSS|Bin)/P(NGSS|Bin)) \div (P(PGSS)/P(NGSS))$$

where NGSS in the Negative Gold Standard Set, P(PGSS|Bin) $\mathrm{P(PGSS|Bin)}$ is the probability that a protein in that bin is in the PGSS, and P(NGSS|Bin)$\mathrm{P(NGSS|Bin)}$ is the probability that a protein in that bin is in the NGSS. $\mathrm{P(PGSS)}$ and $\mathrm{P(NGSS)}$ are the probabilities, independent of bin information, of a protein being in the PGSS and NGSS, respectively. Based on the Spearman correlation coefficient (ρ), PrePPI, MINDy, VIPER and LINCS evidences were weakly correlated, with pairwise ρ varying between -0.06 and 0.11, as required for NB classification. The AP-MS data consists of peptide counts for proteins detected to interact with TBK1, RALA, RALB, and/or RALGDS. However, for proteins that interact with more than one of the bait proteins, the data are highly correlated (ρ > 0.5) across bait proteins. Therefore, in these instances, the maximum of the AP-MS LRs was taken. A final prediction score for each protein's involvement in KRAS regulated signaling is given by multiplying the individual LRs:

$$LR_{Post} = LR_{AP\text{-}MS} * LR_{LINCS} * LR_{DeMAND} * LR_{MINDy} * LR_{VIPER\_upstream} * LR_{VIPER\_downstream} * LR_{PrePPI}$$

where each protein obtains $LR_{Post} LR_{Final}$ as a score for membership in the KRAS PC-Map. Note that the VIPER analysis has two likelihood ratios assigned to it, corresponding to the two analyses described above for upstream regulators and downstream effectors.

Training was performed using two-fold cross validation with holdout, which creates an independent training and testing set and produces a final LR for every protein that is parameterized on the set to which it does not belong. To prioritize novel candidates for experimental validation, the two sets of predictions were rank-transformed separately and combined into one final ranked set. We then selected novel predictions seriatim, skipping those predictions that, based on a literature search, are reported to be involved in KRAS regulated signaling. A list all of predictions is included in **Supplemental File 2**. From the top 40 predicted KRAS pathway proteins, 18 are supported in the literature, and the literature sources are also provided in **Supplemental File 2.** We tested the remaining 22 novel predictions.

To calculate correlated expression between KRAS and other genes, we computed the Pearson correlation coefficient between KRAS expression and all other gene expression in the LUAD gene expression profiles.

**Random Forest Classification**

**Networks using Random Forest Classification**

1. The ARACNe algorithm with adaptive partitioning was used to predict gene regulatory networks for KRAS in the LUAD, LUSC, and COAD contexts with the same parameters as previously described [24]. For LUAD, this resulted in a network of 532,852 TF/CoTF-gene interactions, and a network of 451,969 SP-gene interactions. The corresponding networks using the COAD TCGA samples had 474,452 and 351,554 interactions.
2. CINDy was implemented as previously described[27,62] to predict post-translational modifications of coTFs/TFs by SPs for the LUAD, LUSC, and COAD datasets. CINDy calculates the conditional mutual information among triplets of SP-CoTF/TF-gene and compares the values to those obtained from a null distribution of random TF-gene pairs and randomized expression profiles of SPs. Using a Poisson distribution model, the number of significant triplets is also calculated for random SP/TF pairs. For the CINDy network a corrected p-value threshold of $10^{-8}$ was used, and SP/TF pairs with less than 44 statistically significant gene triplets were discarded. We determined the threshold of 44 triplets for the CINDy network as follows: Based on a previous precision/recall benchmark of CINDy in LUAD[62], we calculated the F1 measure of accuracy (F1 = 2 x (precision x recall)/(precision + recall))[110] as a function of the number of statistically significant triplets. The triplet threshold that corresponded to the

highest F1 score (and thus the highest accuracy) was 44. For LUAD and COAD samples, this resulted in 655,577 and 1,173,102 interactions, respectively.

3. VIPER was applied to LUAD and COAD samples separately. For each tissue type, VIPER predicted the co-segregation between all nsSNPs and the activities for all SPs, CoTFs, and TFs. The p-value of the co-segregation was calculated using aREA [24] and only associations between nsSNP and activity with p-values less than or equal to 0.05 were kept. For LUAD samples, this resulted in a network of 242,695 mutation-activity interactions. For COAD samples, this resulted in 746,447 interactions.
4. Protein-protein interactions from the PrePPI database[28] were retrieved. The $LR_{GO}$ and $LR_{Exp}$ components were removed from $LR_{PrePPI}$, and interactions with modified $LR_{PrePPI}$ scores ≥ 600 were used. This resulted in a total of 732,198 interactions. The PrePPI predictions are the same for both LUAD and COAD contexts.

The ARACNE, VIPER, and CINDy networks for Lung Squamous Cell Carcinoma (LUSC) were derived analogously, using 482 LUSC samples from the TCGA.

**Compilation of PGSS and NGSS for Oncogenic/Tumor Suppressor Pathways**

The PGSS used for training the NB classifier consists of 350 proteins annotated as participating in KRAS signaling pathways by the Ingenuity Pathway Analysis database class "KRAS Molecular Function" (IPA®, QIAGEN Redwood City, www.qiagen.com/ingenuity).

The PGSSs for RF classification were compiled as the union of the KEGG, Biocarta, and Reactome databases from the MSigDB C2 category[65]. In addition, for

KRAS, HIPPO, AMPK and PI3K pathways, the KEGG website[5] was culled for additional pathways that were not recorded in MSigDB C2.

AMPK PGSS: “AMPK Signaling Pathway” from the KEGG database (entry: hsa04152) and MSigDB gene sets: “Reactome Activated AMPK Stimulates Fatty Acid Oxidation in Muscle”, “Reactome Regulation of AMPK Activity Via LKB1”, “Reactome Energy Dependent Regulation of MTOR by LKB1 AMPK”, “Rectome Regulation of RHEB Gtpase activity by AMPK.”

CDKN2A PGSS: “BioCarta ARF Pathway”, “BioCarta CellCycle Pathway” and “KEGG Cell Cycle” gene sets from MSigDB.

EGFR PGSS: “BioCarta EGFR SMRTE pathway” and “KEGG ERBB signaling pathway” gene sets from MSigDB

HIPPO PGSS: “Reactome Signaling by Hippo” from MSigDB and “HIPPO Signaling Pathway” (entry: hsa04390) from the KEGG website.

KRAS PGSS: “RAS Signaling Pathway” from KEGG (entry hsa04014) and the MSigDB gene sets “BioCarta Ras Pathway”, “Reactome Signaling to Ras”, “Reactome Ras activation Upon CA[2] influx through NMDA Receptor”, and “Reactome CREB Phosphorylation through the activation Ras.”

MAPK PGSS: MSigDB gene sets: “BioCarta MAPK Pathway”, “BioCarta P38MAPK Pathway”, “BioCarta Barr MAPK Pathway”, “Reactome P38 MAPK events”, “Reactome Gastrin CREB Signaling Pathway Via PCK and MAPK”, “Reactome ERK MAPK Targets”, “Reactome P130CAS Linkage to MAPK Signaling for Integrins”, “Reactome GRB2:SOS Provides Linkage to MAPK Signaling for Integrins”, “Reactome MAPK Kinase Activation in TLR Cascade”, “Reactome MAPK targets nuclear events mediated by MAP Kinases", "Reactome Activated TAK1 mediates P38 MAPK Activation", "Reactome Traf6 mediated inducation of NFKB and MAPK Kinases upon TLR7 or 9 activation " and "Reactome RAF MAP Kinase Cascade."

NTRK PGSS: "KEGG Neurotrophin Signaling Pathway" gene set from MSigDB.

PI3K PGSS: "Reactome PI3K events in ERBB4 signaling", "Reactome PI3K events in ERBB2 signaling", "Reactome PI3K AKT Activation", "Reactome CD28 dependent PI3K AKT Signaling", "Reactome PI3K Cascade" and the KEGG entry "PI3K-Akt signaling pathway" (entry: hsa04151).

WNT PGSS: MSigDB gene sets "Reactome Signaling by WNT", "KEGG WNT Signaling Pathway" and "BioCarta WNT Pathway."

TP53 PGSS: MSigDB gene sets "BioCarta P53Hypoxia Pathway", "BioCarta P53 Pathway", "Reactome P53 Dependent G1 DNA Damage Response" and "KEGG P53 Signaling Pathway."

The negative gold standard sets (NGSSs) for each pathway are the proteins in the human proteome, as represented by UniProt (n=20,267), with the appropriate PGSS removed.

**Parameters for Random Forests Classifier**

Random Forest classification was implemented using the randomForest[111] R package. The features derived from the networks were as follows: Mutual information for ARACNe, number of statistically significant triplets for CINDy, negative log p-value for VIPER, and LR for PrePPI. We coded each feature symmetrically, so that interactions between protein A and protein B were input into the matrix twice, once in the feature vector for A and once in the feature vector for B. Similar to Havugimana et al.[112], all non-existent interactions and interactions below the respective thresholds were assigned a value of 0 (zero) in the Random Forest feature matrix. Those proteins in the UniProt proteome that had 0s for all features (i.e. did not appear in any of the networks) were removed prior to running the classifier. For each of the 10 oncogene-centric interactomes, proteins that are part of the PGSS were assigned a "1" within the PGSS vector (**Figure 3B, gold column),** while all other proteins were assigned a "0" to represent membership in the NGSS. Training and testing of the Random Forest classifier then proceeded using each pathway's PGSS and NGSS.

**Training and Testing Using Monte-Carlo Cross Validation**

For each oncogene/tumor suppressor, the RF classifier [20] was trained and tested using Monte Carlo cross-validation[105]. Specifically, a forest is trained on equal-sized subsets of the PGSS and NGSS, where the size of the subsest is half of the total PGSS. For the trees in a given forest, the training set is sampled (with replacement) from the PGSS and NGSS subsets. During training, sampled features that best differentiate the PGSS and NGSS samples split the data into successive child nodes, and this continues until all PGSS and NGSS members are separated. The resulting Random Rorest was used to classify the training set (i.e. all proteins excluding the training set) and, thus, predict a probability that each protein is a functional partner of the oncogene in question. The overall procedure occurs 50 times for 50 forests. An OncoSig score of 0.5 means that 50% of trees voted for a protein's association in an oncogenic interactome. All other parameters were set to the default randomForest R package options. RF classification, thus, leverages the consensus of many weak learners while ensuring no crossover between training and testing sets. Critically, using Monte-Carlo cross validation ensures that predictions for each protein are generated with only those random forests that did not include the protein in the training sets. Novel predictions are defined as those that do not appear in a PGSS and have an OncoSig score of at least 0.5.

**Comparing LUAD and COAD results**

To calculate the correlation between LUAD and COAD and between LUAD and LUSC, we performed 100 OncoSig runs each with the LUAD and COAD networks. Distributions of Pearson correlation coefficients were estimated by calculating all pairwise Pearson correlation coefficients between LUAD-LUAD scores, COAD-COAD scores, LUSC-LUSC scores, LUAD-COAD scores, and LUAD-LUSC scores. The statistical significance of the difference, for example, between LUAD-COAD and LUAD-LUAD Pearson correlation coefficients was assessed using Welch's Two Sample t-test.

**Non-redundant PGSS for KRAS**

CD-HIT[104] was used to generate a Non-Redundant KRAS PGSS. The KRAS PGSS was clustered at an 80% sequence identity threshold and, for each cluster, the representative with the longest sequence was selected as per CD-HIT protocol. This resulted in a non-redundant PGSS of 228 proteins in comparison to 250 proteins in the full PGSS. At the 70% and 90% thresholds, the non-redundant PGSSs included 214 and 243 proteins.

**Compilation of Literature-Derived, GO and Ingenuity Sets**

KRAS synthetic lethality and drug-dependency data were compiled from Barbie et al.[54], Corcoran et al.[70], Hayes et al.[71], Astsaturov et al.[73] and GO:0000302: "Response to Reactive Oxygen Species"[6].

When testing for enrichment between the synthetic lethal sets and OncoSig predictions, we created (when possible) a tissue specific network that corresponds to the tissue context for which the original experiment was performed. When this was not possible, we used LUAD-derived networks. Barbie et al. used an integrative method across 19 cell line types (including lung cancer) to detect synthetic lethal KRAS partners, and we thus used LUAD-derived networks. The experiments in Corcoran et al. were done in colorectal cells, and we used COAD-derived networks. The experiments in Hayes et al. were done in pancreatic cell lines, but due to the stroma infiltrate of PDAC tumor samples[113] that would complicate creating an accurate regulatory network, we used LUAD-derived networks for this set. The experiments in Shaw et al. [72], which detected enrichment of proteins involved in "Reactive Oxygen Species Response," were done in mouse epithelial fibroblasts, for which we could not create a network, so we used LUAD-derived networks. The list of proteins involved in EGFR signaling curated by Astsaturov et al. is not context specific, so we used LUAD-derived networks here as well.

To avoid biasing enrichment results to members of the signatures that overlap with the KRAS PGSS, we removed all members of KRAS PGSS that intersected with the respective signatures prior to training. For example, of the 204 synthetic lethal KRAS partners detected by Barbie et al., 23 intersected with the 250 members of KRAS PGSS. Thus, when testing for enrichment with Barbie et al., OncoSig classification was performed on the 227 members of KRAS PGSS that did not overlap, and enrichment was performed with the full 204 synthetic lethal set.

For all of the above literature-derived sets, published protein IDs (usually in Gene Symbol) were converted to UniProt IDs, and those that did not have a UniProt ID were excluded from further analysis.

We used STRINGv10[68] and HumanNet v1[69], when comparing to Oncosig Random Forest results.

**Domain Enrichment**

To test for protein domain enrichment within the KRAS PC-MAP, we downloaded the protein family database PFAM version 31[103]. Families that contained less than 15 proteins were excluded, leaving a total of 351 families. We then performed GSEA using the 351 families and the ranked Random Forest OncoSig scores for the KRAS PC-MAP, excluding the members of the gold standard that was used for training.

**Tandem Affinity Purification**

5 mL packed cell volume of RPE-hTERT cells expressing LAP-tagged proteins were resuspended with 20 mL of LAP-resuspension buffer (300 mM KCl, 50 mM HEPES-KOH [pH7.4], 1 mM EGTA, 1 mM MgCl2, 10% glycerol, 0.5 mM DTT, and protease inhibitors [PI88266, Thermo Scientific]), lysed by gradually adding 600 µL 10% NP-40 to a final concentration of 0.3%, then incubated on ice for 10 min. The lysate was first centrifuged at 14,000 rpm (27,000 g) at 4°C for 10 min, and the resulting supernatant was centrifuged at 43,000 rpm (100,000 g) for 1 hr at 4°C to further clarify the lysate. High speed supernatant was mixed with 500 µL of GFP-coupled beads (Torres et al., 2009) and rotated for 1 hr at 4°C to capture GFP-tagged proteins, and washed five times with 1 mL LAP200N buffer (200 mM KCl, 50 mM HEPES-KOH [pH 7.4], 1 mM EGTA, 1 mM MgCl2, 10% glycerol, 0.5 mM DTT, protease inhibitors, and 0.05% NP40). After re-suspending the beads with 1 mL LAP200N buffer lacking DTT and protease inhibitors, the GFP-tag was cleaved by adding 5 µg of TEV protease and rotating tubes at 4°C overnight. All subsequent steps until the cutting of bands from protein gels were performed in a laminar flow hood. TEV-eluted supernatant was added

to 100 µL of S-protein agarose (69704-3, EMD Millipore) to capture S-tagged protein. After washing three times with LAP200N buffer lacking DTT and twice with LAP100 buffer (100 mM KCl, 50 mM HEPES-KOH [pH 7.4], 1 mM EGTA, 1 mM MgCl2, and 10% glycerol), purified protein complexes were eluted with 50 µL of 2X LDS buffer and boiled at 95°C for 3 min. Samples were then run on Bolt® Bis-Tris Plus Gels (NW04120BOX, Thermo Fisher Scientific) in Bolt® MES SDS Running Buffer (B000202, Thermo Fisher Scientific). Gels were fixed in 100 mL of fixing solution (50% methanol, 10% acetic acid in Optima™ LC/MS grade water [W6-1, Thermo Fisher Scientific]) at room temperature, and stained with Colloidal Blue Staining Kit (LC6025, Thermo Fisher Scientific). After the buffer was replaced with Optima™ water, the bands were cut into eight pieces, followed by washing twice with 500 µL of 50% acetonitrile in Optima™ water. The gel slices were then reduced and alkylated followed by destaining and in-gel digestion using 125 ng Trypsin/LysC (V5072, Promega) as previously described (Shevchenko et al., 2006) with the addition of Protease Max (V2071, Promega) to increase digestion efficiency. Tryptic peptides were extracted from the gel bands and dried in a speed vac. Prior to LC-MS, each sample was reconstituted in 0.1% formic acid, 2% acetonitrile, and water. NanoAcquity (Waters) LC instrument was set at a flow rate of either 300 nL/min or 450 nL/min where mobile phase A was 0.2% formic acid in water and mobile phase B was 0.2% formic acid in acetonitrile. The analytical column was in-house pulled and packed using C18 Reprosil Pur 2.4 uM where the I.D. was 100 uM and the column length was 20-25 cm. Peptide pools were directly injected onto the analytical column in which linear gradients (4-40% B) were of either 80 or 120 min eluting peptides into the mass spectrometer. Either the Orbitrap Elite or Orbitrap Fusion mass spectrometers were used, where a top 15 or "fastest" MS/MS data acquisition was used, respectively. MS/MS was acquired using CID with a collisional energy of 32-35. In a typical analysis, RAW files were processed using Byonic (Protein Metrics) using 12 ppm mass accuracy limits for precursors and 0.4 Da mass accuracy limits for MS/MS spectra. MS/MS data was compared to an NCBI Genbank FASTA database containing all human proteomic isoforms with the exception of the tandem affinity bait construct sequence and common contaminant proteins. Spectral counts were assumed to have undergone fully specific proteolysis and allowing up to two missed cleavages per peptide. All data was filtered and presented at a 1% false discovery rate (Elias and Gygi, 2007). Post processing using in-house MatLab and Excel scripts was used to further process the data for gene ontological analyses.

**Primary Tumor Propagating Cell Culture and Screening Methodology**

Primary lung tumor cells from $KRAS^{G12D/+}$; $p53^{fl/fl}$ mice were cultured in Matrigel as described previously[52]. Prior to seeding, primary cells were infected with a pool of 100-150 lentiviral pLKO shRNAs composed of 3-5 shRNAs per gene at a Multiplicity of Infection <0.5 to ensure single shRNA integration and selected with 1ug/ml puromycin 24 hours after seeding. We screened the top 22 predicted genes from the NB classifier in two pools. Pools also included other candidate vulnerabilities identified by literature review and other methods. 25 pools consisting of 2,286 shRNAs targeting 515 genes not anticipated to be involved in KRAS-regulated signaling were used as a background comparison. After 7 days of spheroid growth, spheroids were dissociated with trypsin into single cells, and half of the 3D culture was re-seeded. The remaining half of each sample was retained for gDNA isolation (T0) until secondary spheroids fully formed 7 days later (T1). The integrated pLKO shRNA was PCR amplified using ExTaq (Clontech), barcoded, multiplexed, and sequenced on an Illumina GAIIx (primer sequences available on request). Sequencing reads were processed into count files in R (v. 3.1.1) using the edgeR package (v. 3.6.8) and analyzed using generalized linear models with edgeR using a time-course design to compare the initial (T0) and final (T1) timepoints and perform a likelihood ratio test[107].

**Calculation of P-values for shRNA Screens and Integration of P-values**

To calculate the statistical significance of the fold change in growth induced by each individual shRNA, we created a density distribution of all the background screens (**Figures 2c and S2, black line**) with area under the curve equal to 1. For each shRNA (in both the KRAS functional partner screen and the BPS screen), we integrated from the minimum $\log_2$FC of the entire BPS to the $\log_2$FC observed for that shRNA, producing a one-tailed p-value for the observed $\log_2$FC. Two shRNAs in the KRAS screen had values below the minimum $\log_2$FC for the BPS, and they were assigned a p-value equivalent to that calculated for the minimum value in the BPS.

**Fisher's Method to Combine shRNA P-values**

Fisher's method can be used to combine p-values that are statistically independent, such as shRNA screens[57,114], and has been used to integrate shRNA KDs previously[114,115]. We used Fisher's method to integrate the p-values of all shRNAs that mapped to the same protein. The formula for calculating a combined p-value given a set of independent p-values is:

$$\chi^2_{2n} = -2 * \sum_{i=1}^{n} \ln(p_i)$$

Where $p_i$ is an individual p-value for an shRNA, $n$ is the total number of shRNAs tested for that protein, and $\chi^2$ is the chi-squared distribution. The integrated p-value is obtained from the chi-squared distribution table with $2n$ degrees of freedom. Using this formula, we integrated all the shRNAs for all 515 genes in the BPS as well as the KRAS functional screen, the positive controls previously reported to be involved in KRAS signaling, and the positive controls reported to be synthetic lethal with $KRAS^{mut.}$. into a single p-value for each gene. For visualization in **Figure 3D**, ties were broken by randomly jittering points along the X axis.

### Gene Enrichment Analysis

Gene enrichment analysis was done by extracting all GO Biological Process terms from the PANTHER database[108], and, for each GO term, testing for overrepresentation between the Naïve Bayes candidates with an integrated p-value<=.05 (red dots to the left of dashed line in **Figure 2D)** and 468 members of BPS (out of 515 total) that were represented in the human Uniprot proteome.

### Multiple Hypothesis Correction

All p-values presented in all analyses were corrected using the Benjamini & Hochberg False Discovery Rate[58] unless noted otherwise.

### Data and Code Availability

**Supplemental File 1** contains the composition of all PGSSs used. A list all of Oncosig-NB predictions is included in **Supplemental File 2**. Raw data and processed files DNA sequencing data are archived in GEO submission GSE107042. Sequences for all RNAi vectors analyzed are provided in **Supplemental File 3**. Code for running Oncosig-NB, Oncosig-RF and perform statistical analysis of the pooled shRNA results are provided in the accompanying codebook.

**ONLINE METHODS REFERENCES:**


109 Biankin, A. V. *et al.* Pancreatic cancer genomes reveal aberrations in axon guidance pathway genes. *Nature* **491**, 399-405, doi:10.1038/nature11547 (2012).

110 Finn, R. D. *et al.* The Pfam protein families database: towards a more sustainable future. *Nucleic Acids Res* **44**, D279-285, doi:10.1093/nar/gkv1344 (2016).

111 Shevchenko, A., Tomas, H., Havlis, J., Olsen, J. V. & Mann, M. In-gel digestion for mass spectrometric characterization of proteins and proteomes. *Nat Protoc* **1**, 2856-2860, doi:10.1038/nprot.2006.468 (2006).

112 Hemming, M. L., Elias, J. E., Gygi, S. P. & Selkoe, D. J. Identification of beta-secretase (BACE1) substrates using quantitative proteomics. *PLoS One* **4**, e8477, doi:10.1371/journal.pone.0008477 (2009).

113 Dai, Z. *et al.* edgeR: a versatile tool for the analysis of shRNA-seq and CRISPR-Cas9 genetic screens. *F1000Res* **3**, 95, doi:10.12688/f1000research.3928.2 (2014).

114 Yu, J., Putcha, P., Califano, A. & Silva, J. M. Pooled shRNA screenings: computational analysis. *Methods Mol Biol* **980**, 371-384, doi:10.1007/978-1-62703-287-2_22 (2013).

115 Tsankov, A. M. *et al.* A qPCR ScoreCard quantifies the differentiation potential of human pluripotent stem cells. *Nat Biotechnol* **33**, 1182-1192, doi:10.1038/nbt.3387 (2015).

116 Mi, H., Poudel, S., Muruganujan, A., Casagrande, J. T. & Thomas, P. D. PANTHER version 10: expanded protein families and functions, and analysis tools. *Nucleic Acids Res* **44**, D336-342, doi:10.1093/nar/gkv1194 (2016).
