## Supplementary Materials for "Systematic Elucidation and Validation of OncoProtein-Centric Molecular Interaction Maps"

### SUPPLEMENTAL INFORMATION

#### Supplemental Note 1:

Gene set enrichment analysis (**Supplemental Table 1**) identifies two related biological processes–i.e., GTPase-mediated signal transduction and intracellular transport/trafficking–as highly enriched in KRAS-specific OncoSig predictions. Additionally, the molecular-layer representation of novel validated interactors (denoted by asterisks in **Figures 2b** and **6**, and italicized below) provides insight into KRAS activity relative to these upstream and downstream processes. In the following paragraphs, we briefly discuss the biology of OncoSig predictions for the KRAS PC-Map.

KRAS regulation of RAB and RHO GTPase signaling remains poorly understood[1,2]. As shown in **Figure 6**, OncoSig predicts both physical and indirect functional interactions between KRAS and (a) RAB family members, (b) RHO family members, and (c) RAB and RHO regulators (GAPs, GEFs, and GDIs), as supported by many previous studies[3-5] (**Figure 2b** and **Figure 6**). Indeed, OncoSig predicts *RAB1A, RAB8A*, *RAB14*, *RAB25,* and *RAB27A*, as well as RAB1B, RAB8B, and RAB32, as downstream effectors and physical binding partners of KRAS. *TBC1D4*, is also a putative RAB GTPase activating protein (GAP)[6] and *APPL1* is an adapter protein that specifically binds to the GTP-bound, active form of RAB5[1]. Interestingly, *RAB1A* has emerged as a novel putative oncogene stimulating tumorigenic growth independent of HRAS signal transduction [7]. Consistently, OncoSig predicted *RAB1A* as a KRAS PC-Map member. Our results are equally intriguing for the class of RHO GTPases. *RHOG, RHOT1*, and CDC42 are all RHO family members predicted as direct KRAS binding partners and downstream effectors (CDC42) by OncoSig. *ARHGDIA*, a RHO GDP-dissociation inhibitor (GDI), has been observed to negatively regulate RHO/RAC signaling[8], *VAV3*[9] and *ARHGEF16*[10] are guanine nucleotide exchange factors (GEFs) for RHO proteins, and *ARHGAP26*[11] is an activator for CDC42[12].

Taken together, our results suggest a far more extensive crosstalk between KRAS signaling and RAB/RHO signaling than previously appreciated[13,14]. They further suggest that KRAS-mediated post-translational regulation of other small-GTPases may be dysregulated in LUAD**.**

Similarly, the role of KRAS in intracellular transport is relatively unknown [15]. Yet, as noted, we have shown compelling connections between KRAS and RAB family members, which are key regulators of intracellular trafficking[2]. In the KRAS PC-Map (**Figure 6**), four RAB family members are inferred as physical KRAS binding partners (bold arrows). In particular, *RAB1A* and *RAB25* mediate ER-Golgi trafficking and transport through apical recycling endosomes, respectively[16], while RAB-interacting proteins *TBC1D4* and *APPL1* promote endosomal vesicular trafficking[1,17]. MAP4Ks, such as *MINK1* (MAP4K6, downstream) and MAP4K1 (upstream) in **Figure 6**, are also implicated in vesicular trafficking through their association with Striatin family complexes, whose dysregulation leads to cancer[18]. Thus, our results suggest that KRAS may be also critically implicated in normal and aberrant intracellular transport.

**Supplemental Figures and Tables**

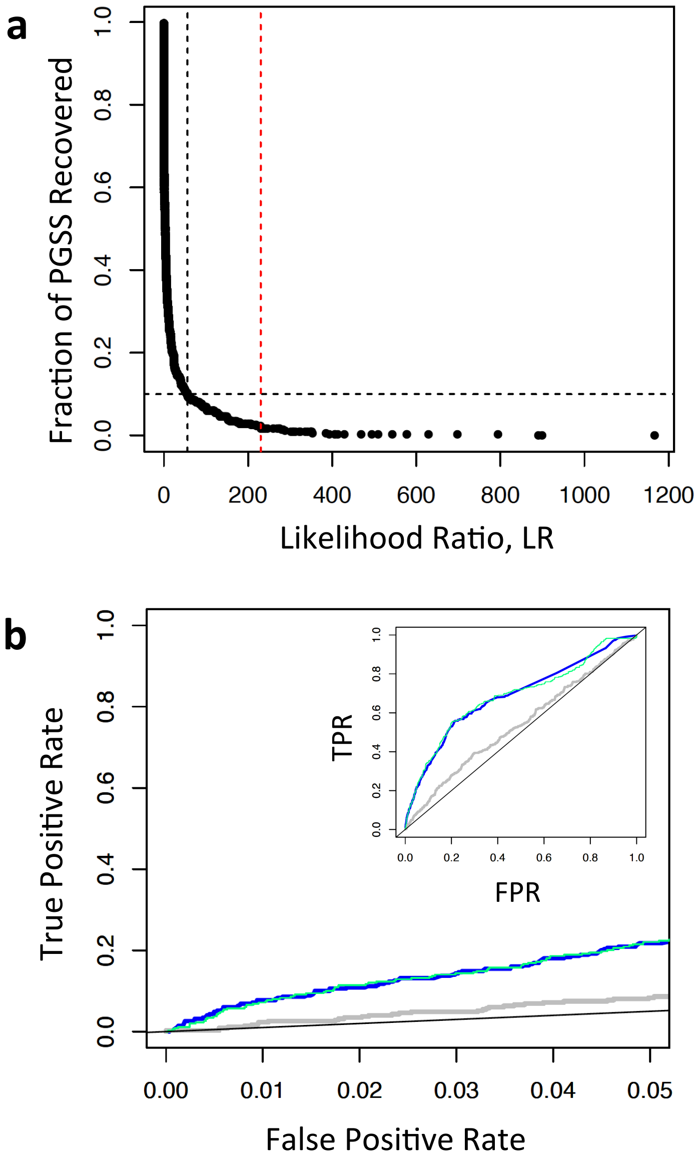


**Figure S1: Recovery of Ingenuity PGSS by the OncoSig Naïve Bayes classifier**

**(a):** Recovery of Ingenuity PGSS (Y axis) as a function of $LR_{\text{Final}}$ (X Axis). At an $LR_{\text{Final}}$ of 56 (vertical black line), the NB classifier recovers 10% of the PGSS (horizontal black line). The vertical red line corresponds to an $LR_{\text{Final}}$ of 230. **(b):** Recovery of Ingenuity PGSS using 2-fold cross-validation (blue), 100-fold Monte-Carlo Cross-validation (teal) and correlated expression (grey).

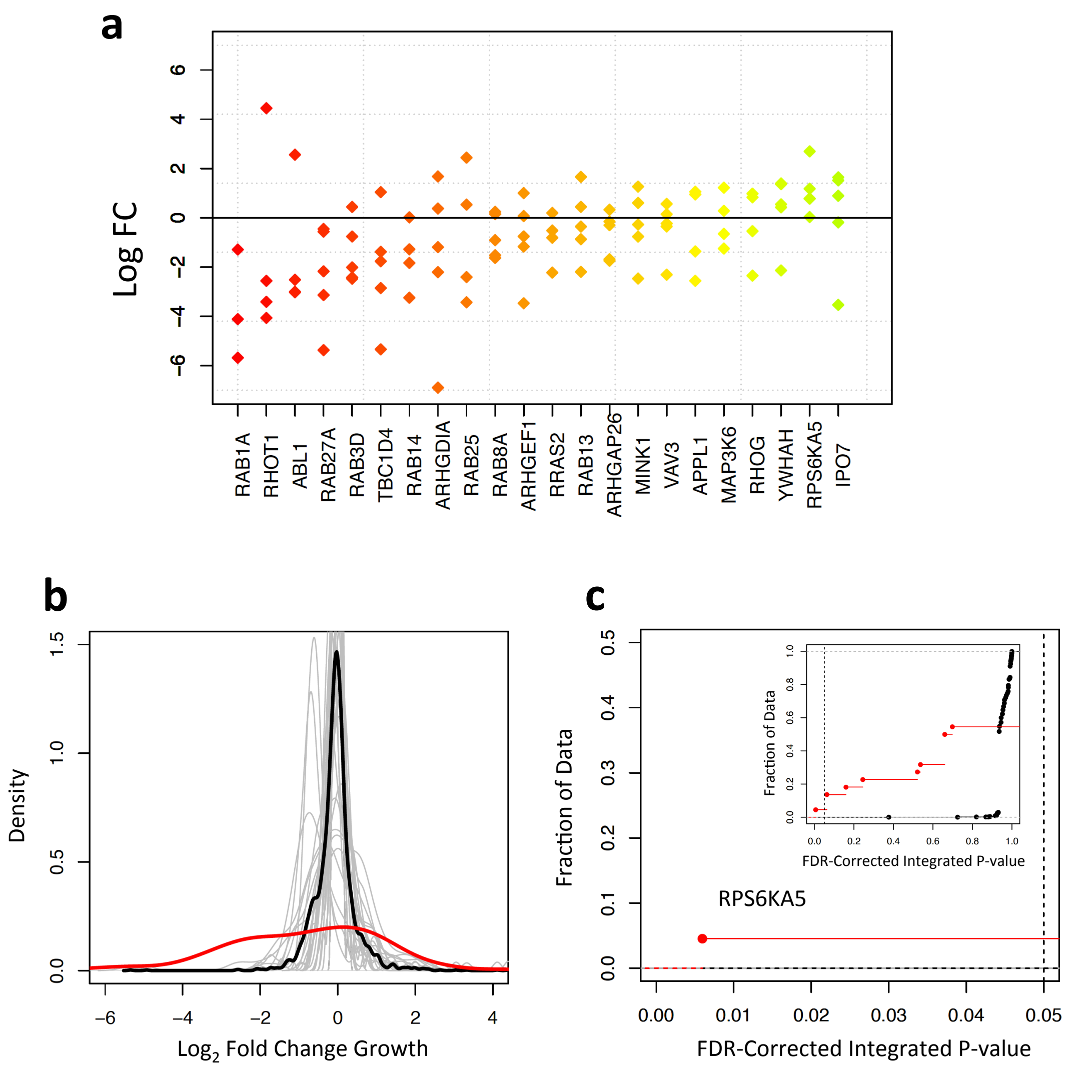


**Figure S2: Screen results for predicted novel KRAS functional partners**

**(a):** The $Log_2FC$ of shRNA abundance is plotted against the novel proteins tested in the KRAS negative selection screen. The 3-5 points plotted for a given protein are shRNAs that target the mRNA for that protein. The X axis is sorted by mean $Log_2FC$ for all shRNAs targeting each gene. Colors change from red to green with mean $Log_2FC$.

**(b):** Density plots of $Log_2FC$ for predicted KRAS functional partners (red), all individual BPS ( grey), and the average of all BPS (black).

**(c):** An empirical cumulative distribution function (eCDF) plot of the corrected integrated p-values for the predicted KRAS functional partners (red) and the 515 proteins in the BPS (black), for increasing cell growth. None of the BPS proteins had a p-value less than .05.

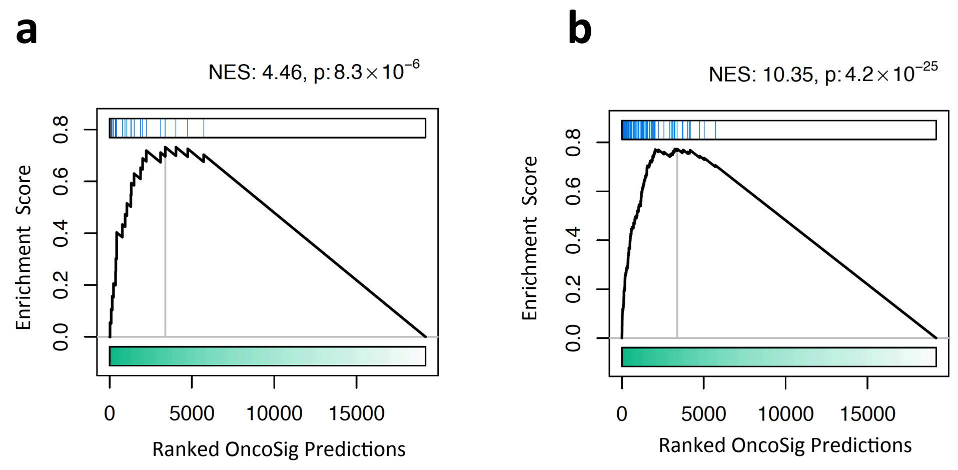


**Figure S3: Comparison of the NB and RF OC-Maps in recovery of Ingenuity Gold Standard Set proteins**

**(a):** GSEA of the 22 predicted KRAS functional partners tested in knockdown experiments where the ranking is based on OncoSig using the KRAS Ingenuity PGGS.

**(b):** Enrichment of the top 100 NB predictions within the OncoSig where the ranking is based on OncoSig using the KRAS Ingenuity PGGS.

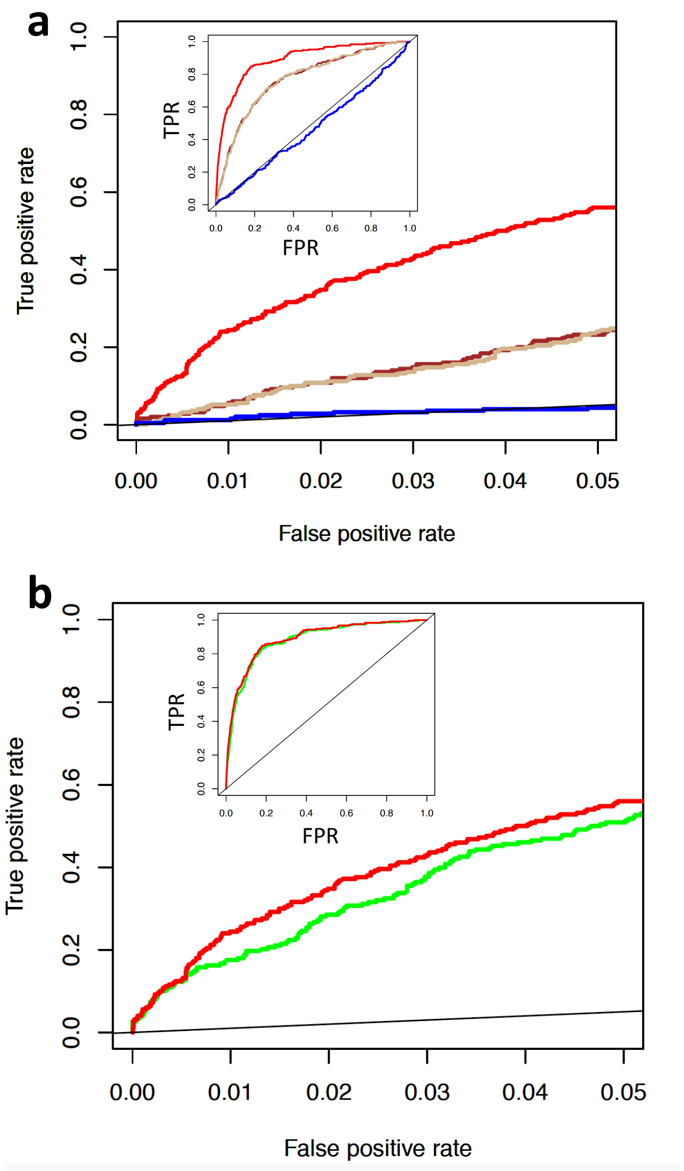


**Figure S4: OncoSig performance with a scrambled network and non-redundant positive gold standard proteins**

**(a):** ROC curves showing performance of node degree ranking (brown),OncoSig trained using a degree-preserving edge shuffled network (tan), and Pearson's correlation coefficient of mRNA expression level with KRAS in LUAD gene expression profiles (blue) in capturing the KRAS PGSS compared to OncoSig trained using a non-scrambled network results (red).

**(b):** Performance of OncoSig using the KRAS non-redundant PGSS (green) and the full PGSS (red). In these plots the red KRAS PGSS ROC curve is the same as the red KRAS PGSS ROC curve in **Figure 3**.

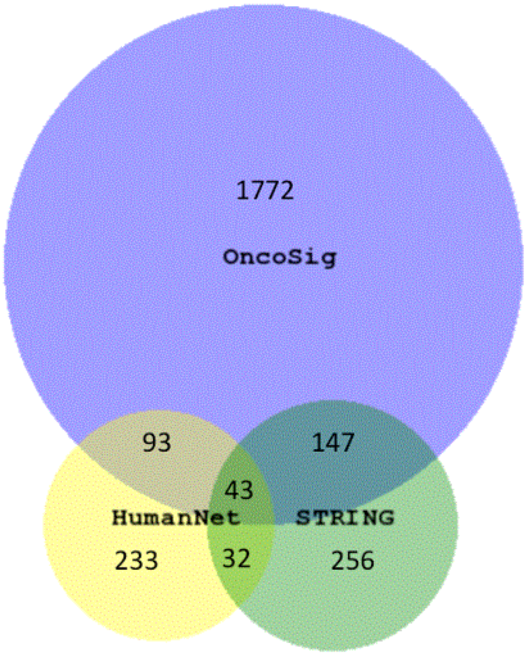


**Figure S5: Comparison of OncoSig and other protein-protein interaction resources**

A Venn diagram showing overlap of OncoSig predictions (score ≥ 0.5 or probability threshold ≥ 50%) trained on KRAS PGSS using LUAD derived networks (blue), and proteins annotated by STRING (score threshold ≥ 400) (green) and HumanNet (Log likelihood threshold ≥ 0) (yellow) as engaging in functional interactions with KRAS.

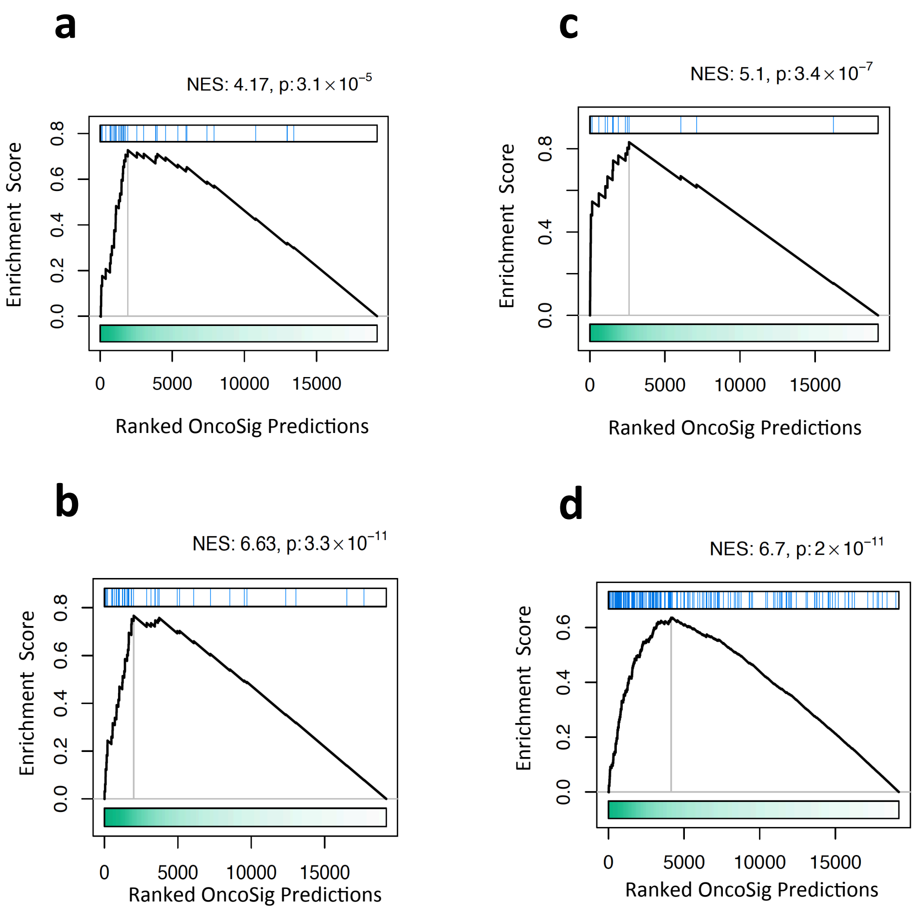


**Figure S6: GSEA of OncoSig results with literature-derived sets**

**(a):** Enrichment of proteins identified as synthetic lethal with mutant KRAS using at least 2 shRNA hairpins or hairpin ranking analysis (Barbie et al., 2009).

**(b):** Enrichment of proteins synthetic lethal to KRAS in the presence of a MEK inhibitor (Corcoran et al., 2013).

**(c):** Enrichment of protein resistance-signature to ERK inhibitor SCH772984 (Hayes et al., 2016).

**(d):** Enrichment of proteins involved in response to Reactive Oxygen Species (GO:0000302) (Ashburner et al., 2000).

Enrichment of sets represented in panels A,C and D are based on OncoSig results from LUAD-derived networks, while enrichment shown in panel B is based on COAD-derived networks.

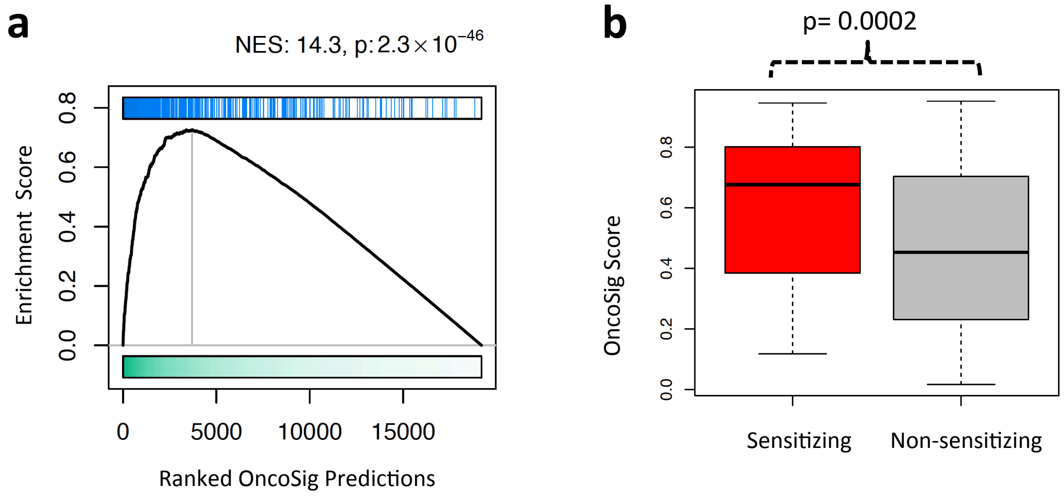


**Figure S7: OncoSig predictions for EGFR are enriched for EGFR signaling and synthetic lethal partners**

**(a)**: Enrichment of literature derived EGFR signaling network (Astsaturov et al., 2010) with OncoSig predictions.

**(b):** Box plots of the OncoSig scores for EGFR synthetic lethal partners (red) and the EGFR curated pathway proteins not identified as synthetic lethal (grey). The p-value was calculated using Welch's two sample t-test.

| GO Term | OncoSig Naïve Bayes Fraction | Background Fraction | Fold Enrichment | p-value (FDR-corrected) |
|---|---|---|---|---|
| regulation of vesicle-mediated transport (GO:0060627) | 0.3889 | 0.0223 | 17.4067 | 0.0004 |
| small GTPase mediated signal transduction (GO:0007264) | 0.3333 | 0.0138 | 24.1345 | 0.0004 |
| regulation of small GTPase mediated signal transduction (GO:0051056) | 0.3333 | 0.0148 | 22.4914 | 0.0004 |
| regulation of localization (GO:0032879) | 0.7222 | 0.1217 | 5.9359 | 0.0021 |
| membrane organization (GO:0061024) | 0.4444 | 0.0452 | 9.8331 | 0.0021 |
| cellular response to insulin stimulus (GO:0032869) | 0.2222 | 0.0071 | 31.0844 | 0.0031 |
| Ras protein signal transduction (GO:0007265) | 0.2222 | 0.0076 | 29.1477 | 0.0034 |
| vesicle-mediated transport (GO:0016192) | 0.5556 | 0.0846 | 6.5689 | 0.0035 |
| establishment of protein localization (GO:0045184) | 0.5 | 0.0684 | 7.3063 | 0.0035 |
| regulation of transport (GO:0051049) | 0.5556 | 0.0882 | 6.2984 | 0.004 |
| Rho protein signal transduction (GO:0007266) | 0.1667 | 0.003 | 54.5363 | 0.004 |
| intracellular signal transduction (GO:0035556) | 0.5 | 0.0772 | 6.477 | 0.0053 |
| positive regulation of catalytic activity (GO:0043085) | 0.5 | 0.0784 | 6.3786 | 0.0053 |
| positive regulation of GTPase activity (GO:0043547) | 0.3333 | 0.0315 | 10.5847 | 0.0053 |
| actin cytoskeleton organization (GO:0030036) | 0.2778 | 0.02 | 13.8559 | 0.0053 |
| response to insulin (GO:0032868) | 0.2222 | 0.0102 | 21.8011 | 0.0053 |
| single-organism organelle organization (GO:1902589) | 0.4444 | 0.0642 | 6.9178 | 0.0061 |
| protein transport (GO:0015031) | 0.4444 | 0.0647 | 6.872 | 0.0061 |
| regulation of GTPase activity (GO:0043087) | 0.3333 | 0.0342 | 9.7472 | 0.0061 |
| peptide transport (GO:0015833) | 0.4444 | 0.0659 | 6.748 | 0.0062 |

**Supplemental Table 1: GO Biological Process Enrichment of Predicted KRAS Functional Partners**
Top 20 enriched GO Biological Processes of the 18 statistically significant (pvalue <=.05) predicted KRAS functional partners (see text and Figure 3C) and all proteins in the human proteome, as defined by Uniprot. Column 1 is the GO Biological Process. Columns 2 and 3 are, respectively, the fraction of predicted KRAS functional partners and the human proteome annotated as part of the Biological Process. Column 4 is the fold enrichment obtained by dividing column 2 by column 3. Column 5 is the False Discovery Rate (FDR) corrected p-values of the fold enrichment.
Biological Processes are sorted by FDR corrected p-values (Column 5).

### REFERENCES


1 Liu, Z., Xiao, T., Peng, X., Li, G. & Hu, F. APPLs: More than just adiponectin receptor binding proteins. *Cellular signalling* **32**, 76-84, doi:10.1016/j.cellsig.2017.01.018 (2017).

2 Tzeng, H. T. & Wang, Y. C. Rab-mediated vesicle trafficking in cancer. *J Biomed Sci* **23**, 70, doi:10.1186/s12929-016-0287-7 (2016).

3 Chew, T. W. *et al.* Crosstalk of Ras and Rho: activation of RhoA abates Kras-induced liver tumorigenesis in transgenic zebrafish models. *Oncogene* **33**, 2717-2727, doi:10.1038/onc.2013.240 (2014).

4 Patel, M. & Cote, J. F. Ras GTPases' interaction with effector domains: Breaking the families' barrier. *Communicative & integrative biology* **6**, e24298, doi:10.4161/cib.24298 (2013).

5 Fritsch, R. *et al.* RAS and RHO families of GTPases directly regulate distinct phosphoinositide 3-kinase isoforms. *Cell* **153**, 1050-1063, doi:10.1016/j.cell.2013.04.031 (2013).

6 Gabernet-Castello, C., O'Reilly, A. J., Dacks, J. B. & Field, M. C. Evolution of Tre-2/Bub2/Cdc16 (TBC) Rab GTPase-activating proteins. *Mol Biol Cell* **24**, 1574-1583, doi:10.1091/mbc.E12-07-0557 (2013).

7 Thomas, J. D. *et al.* Rab1A is an mTORC1 activator and a colorectal oncogene. *Cancer Cell* **26**, 754-769, doi:10.1016/j.ccell.2014.09.008 (2014).

8 Lu, W. *et al.* Downregulation of ARHGDIA contributes to human glioma progression through activation of Rho GTPase signaling pathway. *Tumour Biol*, doi:10.1007/s13277-016-5374-6 (2016).

9 Hornstein, I., Alcover, A. & Katzav, S. Vav proteins, masters of the world of cytoskeleton organization. *Cellular signalling* **16**, 1-11 (2004).

10 Oliver, A. W. *et al.* The HPV16 E6 binding protein Tip-1 interacts with ARHGEF16, which activates Cdc42. *Br J Cancer* **104**, 324-331, doi:10.1038/sj.bjc.6606026 (2011).

11 Burroughs, A. M., Balaji, S., Iyer, L. M. & Aravind, L. Small but versatile: the extraordinary functional and structural diversity of the beta-grasp fold. *Biology direct* **2**, 18, doi:10.1186/1745-6150-2-18 (2007).

12 Doherty, J. T. *et al.* Skeletal muscle differentiation and fusion are regulated by the BAR-containing Rho-GTPase-activating protein (Rho-GAP), GRAF1. *The Journal of biological chemistry* **286**, 25903-25921, doi:10.1074/jbc.M111.243030 (2011).

13 Boulter, E., Estrach, S., Garcia-Mata, R. & Feral, C. C. Off the beaten paths: alternative and crosstalk regulation of Rho GTPases. *Faseb J* **26**, 469-479, doi:10.1096/fj.11-192252 (2012).

14 Cox, A. D. & Der, C. J. Ras history: The saga continues. *Small GTPases* **1**, 2-27, doi:10.4161/sgtp.1.1.12178 (2010).

15 Prior, I. A. & Hancock, J. F. Ras trafficking, localization and compartmentalized signalling. *Seminars in cell & developmental biology* **23**, 145-153, doi:10.1016/j.semcdb.2011.09.002 (2012).

16 Bhuin, T. & Roy, J. K. Rab proteins: the key regulators of intracellular vesicle transport. *Exp Cell Res* **328**, 1-19, doi:10.1016/j.yexcr.2014.07.027 (2014).

17 Fukuda, M. TBC proteins: GAPs for mammalian small GTPase Rab? *Biosci Rep* **31**, 159-168, doi:10.1042/BSR20100112 (2011).

18 Hwang, J. & Pallas, D. C. STRIPAK complexes: structure, biological function, and involvement in human diseases. *Int J Biochem Cell Biol* **47**, 118-148, doi:10.1016/j.biocel.2013.11.021 (2014).
